# Firing patterns of serotonin neurons underlying cognitive flexibility

**DOI:** 10.1101/059758

**Authors:** Sara Matias, Eran Lottem, Guillaume P. Dugué, Zachary F. Mainen

**Affiliations:** Champalimaud Research, Champalimaud Centre for the Unknown, Avenida Brasilia, s/n, 1400-038, Lisbon, Portugal; MIT-Portugal Program; Institut de Biologie de l’Ecole Normale Supérieure, Centre National de la Recherche Scientifique (CNRS), UMR8197, Institut National de la Santé et de la Recherche Mdicale (INSERM) U1024, Paris, France

## Abstract

Serotonin is implicated in mood and affective disorders^1,2^ but growing evidence suggests that its core endogenous role may be to promote flexible adaptation to changes in the causal structure of the environment^3–8^. This stems from two functions of endogenous serotonin activation: inhibiting learned responses that are not currently adaptive^9,10^ and driving plasticity to reconfigure them^11–13^. These mirror dual functions of dopamine in invigorating reward-related responses and promoting plasticity that reinforces new ones^16,17^. However, while dopamine neurons are known to be activated by reward prediction errors^18,19^, consistent with theories of reinforcement learning, the reported firing patterns of serotonin neurons^21–23^ do not accord with any existing theories^1,24,25^. Here, we used long-term photometric recordings in mice to study a genetically-defined population of dorsal raphe serotonin neurons whose activity we could link to normal reversal learning. We found that these neurons are activated by both positive and negative prediction errors, thus reporting the kind of surprise signal proposed to promote learning in conditions of uncertainty^26,27^. Furthermore, by comparing cue responses of serotonin and dopamine neurons we found differences in learning rates that could explain the importance of serotonin in inhibiting perseverative responding. Together, these findings show how the firing patterns of serotonin neurons support a role in cognitive flexibility and suggest a revised model of dopamine-serotonin opponency with potential clinical implications.

---

We investigated serotonin (5-HT) function in a reversal learning task in which the associations between neutral odour cues and different positive and negative outcomes are first well learned and then suddenly changed. We reasoned that the scarcity of prediction error like responses in previous recordings of identified 5-HT neurons^21–23^ or unidentified raphe neurons^28,29^ might be due to inadequately strong prediction errors, which could be much stronger in a reversal task. We first sought causal evidence that 5-HT neurons were linked to reversal learning in this task by using a pharmacogenetic approach to silence 5-HT neurons^30–32^. Transgenic mice expressing CRE recombinase under the 5-HT transporter promoter^33^ (SERT-Cre, n=8) were transduced with a Cre-dependent adeno-associated virus (AAV.Flex) expressing the synthetic receptor Di (DREADD, hM4D)^32^ injected in the dorsal raphe nucleus (DRN), the major source of 5-HT to the forebrain (Fig. 1a). These mice and their wild type littermates (WT, n=4) were trained in a head-fixed classical conditioning paradigm in which one of four odour cues (conditioned stimuli, CS’s) was randomly presented in each trial. After a fixed 2 s trace period, each odour was followed by a tone and a specific outcome, or unconditioned stimulus (US) (Fig. 1b top). For two odours the US was a water reward and for the other two it was nothing (i.e. only the tone was played). After training, mice showed learning of the odour-outcome contingencies, as indicated by differences in anticipatory lick rate (Fig. 1b bottom).

To test the impact of inhibiting DRN SERT-Cre expressing neurons (hereafter simply “5-HT neurons”) we used a within-animal cross-over design in which each mouse experienced two reversals (Fig. 1c top), receiving the DREADD ligand clozapine-*N*-oxide (CNO) during one and vehicle during the other; WT mice, which always received CNO, served as additional controls (Extended Data Fig. 1a). As expected, mice adjusted their anticipatory licking according to the reversed associations in both reversals (Fig. 1c bottom, grey traces). For worse than expected outcomes (negative reversals), the kinetics of adaptation to the new contingencies was significantly slower in hM4D mice receiving CNO compared to hM4D controls and WT controls (Fig. 1c bottom red and black traces, 1d; Extended Data Fig. 1b). In contrast, for better than expected outcomes (positive reversals), there was no difference between treatment and control groups (Fig. 1d).

This experiment shows that a specific population of 5-HT neurons in the DRN contribute to inhibiting perseverative responding, suggesting an anatomical and genetic substrate for previous results obtained with pharmacological and lesion experiments^3–7^. These findings also defined a genetic access point to assess how the net activity of a specific population of 5-HT neurons could account for its effects on reversal learning. For this, we used photometry to monitor the activity of these DRN 5-HT neurons through an implanted optical fibre^34^ (Fig. 2a). SERT-Cre mice were infected in the DRN using two AAV.Flex viruses containing the genetically-encoded calcium indicator GCaMP6s^35^ and the activity-insensitive fluorophore, tdTomato (Fig. 2b). We verified the specificity of GCaMP6s expression to DRN 5-HT neurons using histological methods (Extended Data Fig. 2a-c). We used a regression-based method to decompose the dual fluorescence signals into a GCaMP6s-specific component, reflecting activity-dependent changes, and a shared component, reflecting general fluorescence changes (e.g. movement artefacts; see Methods, Extended Data Fig. 3). We validated the effectiveness of this approach in control mice (n=4) infected in the DRN with yellow fluorescent protein (YFP; replacing GCaMP) and tdTomato (Extended Data Figs. 2d-h, Fig 3).

**Figure 1.**
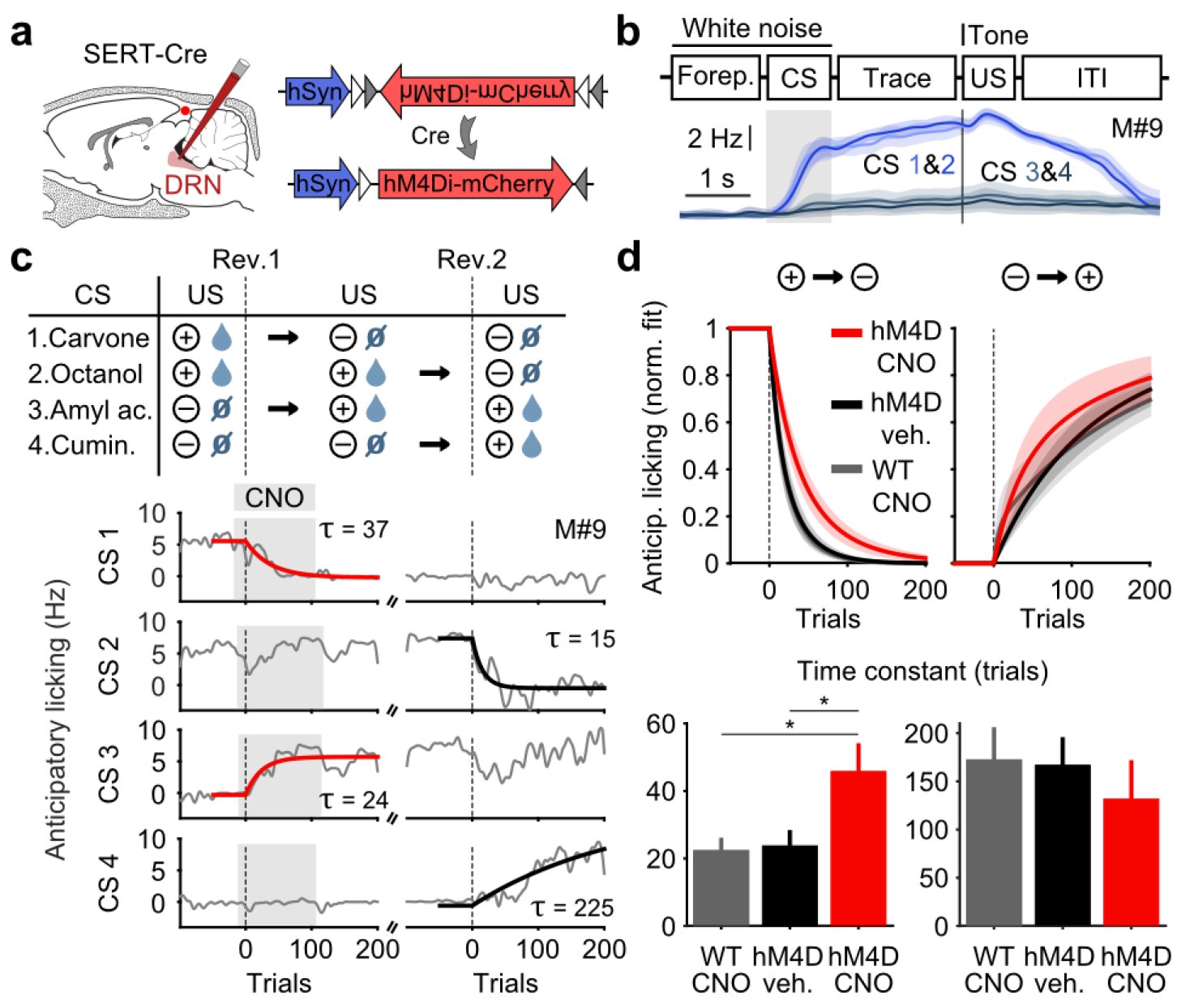
Inhibition of DRN 5-HT neurons causes perseverative responding. **a**, Injections of Cre-dependent hM4Di-mCherry (right) in the DRN of SERT-Cre mice (left). **b**, Trial structure of the task (top) and mean lick rate of an example session along the four trial types (bottom). **c**, Reversals procedure (top) and example of adaptation in mean anticipatory licking across trials around reversals (bottom, grey) with exponential fits to reversed odours (red and black traces). Grey shade represents trials of sessions after CNO injection. **d**, Mean exponential fits of anticipatory licking for each group of mice after reversal (top) and corresponding mean time constants (bottom, 1way ANOVA, F_2,19_ = 6.28, *p* = 0.008 for negative reversal, F_2,16_ = 0.34, *p* = 0.715 for positive reversal; multiple comparisons signalled in the figure). \**p* < 0.05.

To obtain a broader picture of DRN 5-HT activity and compare our results to other DRN recording studies^22,29^, mice learned to associate four odours with four different outcomes: large water reward, small water reward, nothing (neutral) and mild air puff to the eye (Extended Data Fig. 4). After approximately two weeks of training mice showed robust CS-triggered anticipatory licking correlated to the reward value of the associated US’s (big water > small water > neutral ≈ air puff, Extended Data Fig. 4). Photometric 5-HT responses were similar to previous electrical^21,22^ and photometric^23^ recordings of identified 5-HT neurons (Fig. 2c, Extended Data Fig. 5). YFP control mice implanted and recorded in the same manner showed no photometric responses (Extended Data Fig. 6). We then reversed the CS-US associations in pairs, such that the CS’s associated with the large and small rewards now predicted the air puff and neutral outcomes respectively, and vice versa (Fig. 2d). Upon this reversal, mice experienced strong violations of CS-based expectations, both positive and negative in value, when the unexpected US’s were delivered. Anticipatory licking measurements showed that mice adapted to reversal of contingencies over 1-3 additional sessions (Extended Data Fig. 7).

To understand the pattern of 5-HT neural activity that could underlie adaptation to reversal of contingencies we first analysed US responses, which could contribute to or modulate reinforcement learning. We found that the abrupt reversal of cue-outcome associations caused immediate changes in 5-HT US responses (Extended Data Fig. 8), including a robust positive response to the absence of a predicted reward (Fig. 2e). The results for all US’s are summarized in Fig. 2f. First, 5-HT neurons showed little or no response to expected water rewards before reversal, but responded robustly to the same rewards when they were unexpected, after reversal. In other words, they reported a positive reward prediction error (RPE) by an increase in firing.

**Figure 2.**
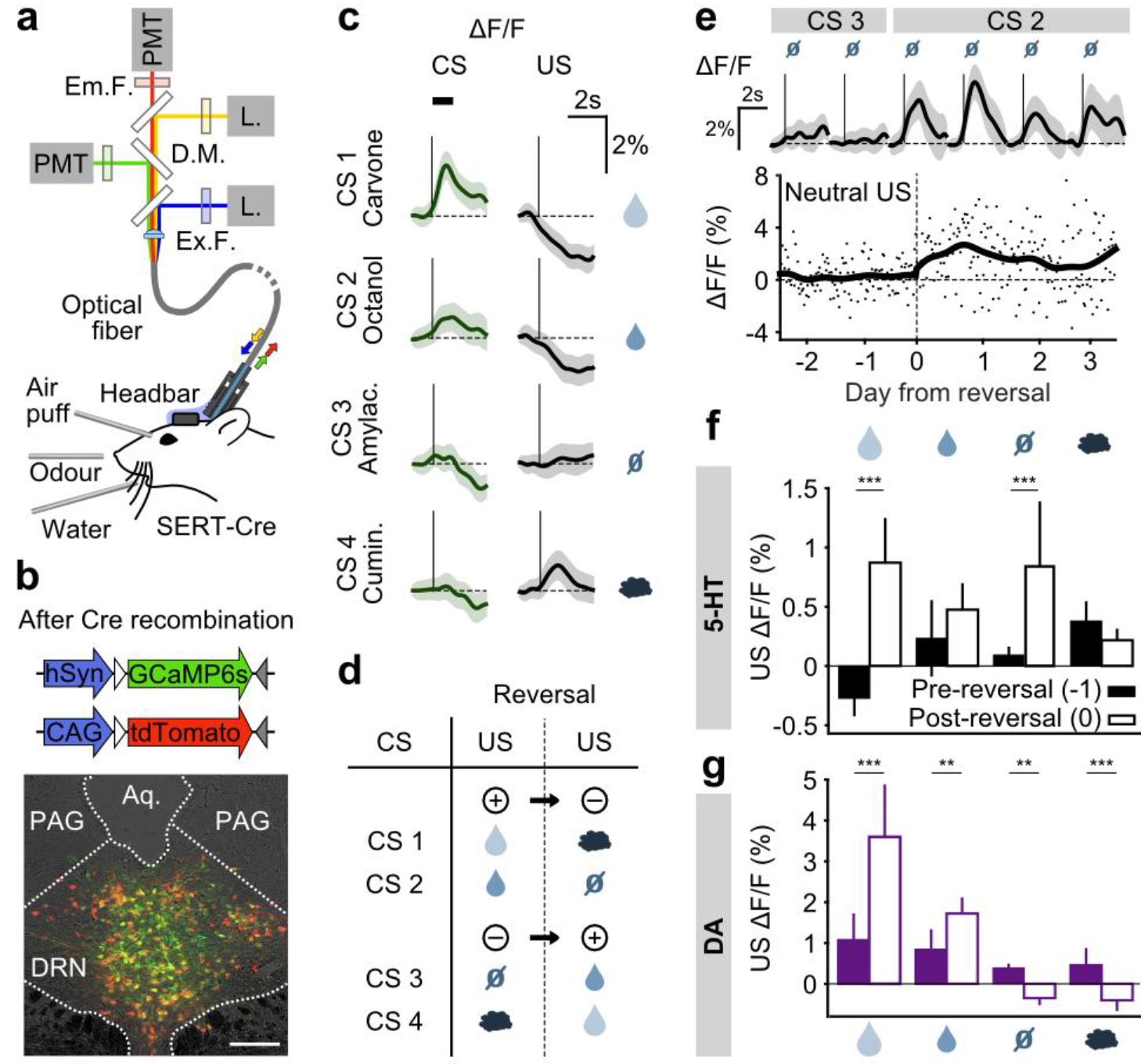
DRN 5-HT neurons signal an unsigned prediction error. **a** Fibre photometry with movement artefact correction in head-fixed mice. L.: laser, PMT: photomultiplier tube, D.M.: dichroic mirror, Ex.: excitation, Em.:emission, F.: filter. **b**, Cre-dependent fluorophores used (top) and coronal section showing expression of GCaMP6s and tdTomato in the DRN of a SERT-Cre mouse (bottom, scale bar: 200 μm). **c**, Mean responses of 5-HT neurons to the four CS’s and US’s during an example session of a mouse before reversal. Shaded areas represent 95% confidence interval (CI). **d**, Reversal procedure (negative reversal: CS 1 and 2, positive reversal: CS 3 and 4). **e** Mean neutral US responses of an example mouse over days around reversal (top, shaded areas represent 95% CI) and change in mean neutral US response amplitude across those days (trace; dots represent individual trials). **f** Mean (± s.e.m.) response of 5-HT neurons across mice to the four US’s before (day-1, filled bars) and right after (day 0, open bars) reversal (n = 8 mice, 2-way ANOVA with factors mouse and day, main effect of day F_1.764_ = 84.36 *p* < 0.001 for large reward, F_1.748_ = 3.49 *p* = 0.062 for small reward, F_1.756_ = 38.17 *p* < 0.001 for neutral, F_1.766_ = 2.79 *p* = 0.095 for air puff). **g** Same as **f** for midbrain DA neurons (n = 3 mice, F_1.249_ = 67.9 *p* < 0.001 for large reward, F_1.277_ = 8.49 *p* = 0.004 for small reward, F_1.278_ = 10.95 *p* = 0.001 for neutral, F_1 250_ = 12.74 *p* < 0.001 for air puff. ** *p* < 0.01, *** *p* < 0.001.

Second, the same 5-HT neurons also showed little response to a neutral US when it was expected but a robust response when it occurred in place of a reward after reversal (Fig. 2e). That is, 5-HT neurons also reported a negative RPE by an increase in firing. Thus, with the strong violations of expectations that occur in a reversal task, 5-HT neurons were found to be very sensitive to prediction errors, much more so than in simple reward omission tests^22,23,28,29^. Excitatory responses to both negative and positive surprises indicate an unsigned RPE, sometimes known as a ‘surprise’ or ‘salience’ signal, as had been theorized for other neuromodulators^36,37^ rather than a signed negative RPE as had been theorized for 5-HT^1,24,25^.

To compare directly how dopamine (DA) neurons respond in the same paradigm, we infected TH-Cre mice and targeted neurons in either the posterior lateral ventral tegmental nucleus (VTA) or the substatia nigra pars compacta (SNc) (Extended Data Fig. 9a). DA photometry responses in these two areas were comparable and were therefore combined (Extended Data Figs. 9b,c; Fig. 10). Like 5-HT neurons, DA neurons showed much stronger excitatory responses to water rewards immediately after reversal when they violated cue-based predictions than before reversal when they occurred as predicted. In contrast to 5-HT neurons, DA neurons showed an inhibitory response to neutral US’s when a water reward was expected (Fig. 2g; Extended Data Fig. 10). Thus, the TH-positive midbrain DA neurons we accessed using fibre photometry were sensitive to RPE’s and could be described as a signed RPE, as reported previously in reward-omission paradigms^18,19^ (but see ^38–40^), and in contrast to the unsigned RPE seen for SERT-positive DRN 5- HT neurons.

To further investigate the idea that 5-HT neurons report a surprise signal, five days after reversal a randomly-selected US was delivered on a small fraction (20%) of trials at the time that a CS was normally presented (Fig. 3a). Water rewards produced larger 5-HT responses when they were un-cued compared to when preceded by a well-learned cue (Fig. 3b). Of particular interest, even neutral tones also produced an excitatory response when an odour was expected (Extended Data Fig. 11a). Therefore, 5-HT neurons are activated by the substitution of one neutral stimulus with another. In this paradigm, DA neurons also responded strongly to un-cued rewards as previously reported^18,19^ but little to other un-cued US’s (Fig.3c).

**Figure 3.**
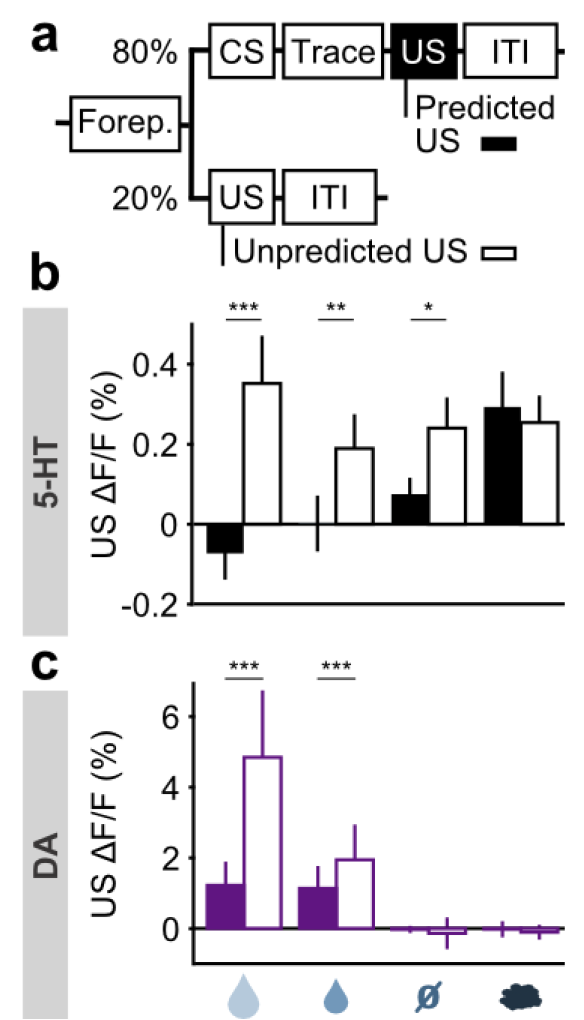
DRN 5-HT neurons report a surprise signal. **a**, Behavioural task including surprise US trials. **b**, Mean (± s.e.m.) response of DRN 5-HT neurons across mice to the four US’s when they are predicted (filled bars) and when they are unpredicted (open bars) (n = 4 mice, 2-way ANOVA with factors type (predicted or unpredicted) and mouse, main effect of type: large reward F_1.923_ = 45.17, *p* < 0.001, small reward F_1.944_ = 8.42, *p* = 0.0038, neutral F_1.924_ = 5.36, *p* = 0.0208, air puff F_1.924_ = 0.61, *p* = 0.4331). **c**, Same as **b** but for midbrain DA neurons (n = 3 mice, large reward F_1.642_ = 175.05, *p* < 0.001, small reward F_1.589_ = 17.53, *p* < 0.001, neutral F_1.673_ = 0.52, *p* = 0.4707, air puff F_1.601_ = 0.34, *p* = 0.5598). \**p* < 0.05, \*\**p* < 0.01, \*\*\**p* < 0.001.

Thus, consistent with the responses following CS-US reversal, this experiment also showed that 5-HT neurons respond in the same manner to surprises whether negative, neutral or positive, whereas DA neurons are primarily sensitive to positive surprises. These finding indicate that 5- HT and DA neurons are both sensitive to violations of expectation that occur during changes in the learned structure of the environment. The two systems responded in the same way to better-than-expected US’s but in opposite ways to worse-than-expected US’s. The strong excitatory response of the 5-HT system to negative RPE’s provides a plausible explanation for why inhibiting this system impairs negative reversal learning^4,6^ (Fig. 1). Unlike signed prediction error signals, surprise or unsigned prediction error signals are not useful teaching signals. But surprise signals are well-suited to represent uncertainty and are therefore useful to increase learning rates or plasticity^26,27^. Thus, during extinction learning or negative reversals, the 5-HT system, either directly or through an interaction with the DA system, could facilitate trial-by-trial un-doing of DA-dependent learning, as suggested by in vitro experiments. 5-HT neurons also respond during positive surprises, such as during positive reversal or initial learning. Under the same account, activation might compete with co-occurring DA signals, which can actually slow positive learning^41^. A second role for surprise signals, aside from enhancing learning, is to reduce the influence of the systems generating the errors^36,37^. 5-HT could thus contribute not only through learning and plasticity but by directly suppressing activity in systems responsible for violated predictions. Indeed, 5-HT has been strongly associated with suppressing both impulsive and perseverative responses through “behavioural inhibition”^1,4,25^. This mirrors and opposes the DA system, which is known to invigorate reward-directed actions^1,15,25^. US surprise responses occur too late in the trial to inhibit maladaptive CS-driven behavioural responses; to intervene in time to prevent responding, behavioural inhibition should be triggered by predictive CS cues. We therefore examined the CS responses of 5-HT and DA neurons. Before reversal (day -1), both showed CS responses that reflected the relative value of the US predicted by the CS (big reward > small reward > neutral ~ air puff) (Extended Data Figs. 5a, Fig. 8b). After reversal, both adjusted to the new contingencies (Extended Data Figs. 12, Fig 13) such that, by day 3 post-reversal, the CS responses reflected their new US associations (Fig. 4a, Fig 4c; open bars). Thus, despite small differences in the relative magnitudes, DA and 5-HT neurons showed CS responses that were similar both before and after reversal learning, in contrast to their distinct US responses.

Since DA and 5-HT are believed to have opposing direct effects on behaviour vigour^1^, the similarity of their CS responses suggests that in steady-state they would tend to cancel each other out. However, when we analysed the time course of the adaptation of these CS responses, we found that 5-HT CS responses had a markedly slower rate of adaptation to the new contingencies than did DA CS responses (Fig. 4b, Fig 4d,Fig. 4e,Fig. 4f). The difference in the time consstant of CS adaptation was significant for both negative and positive reversals and was not due to differences in learning rates between groups of mice (Fig. 4e).

**Figure 4.**
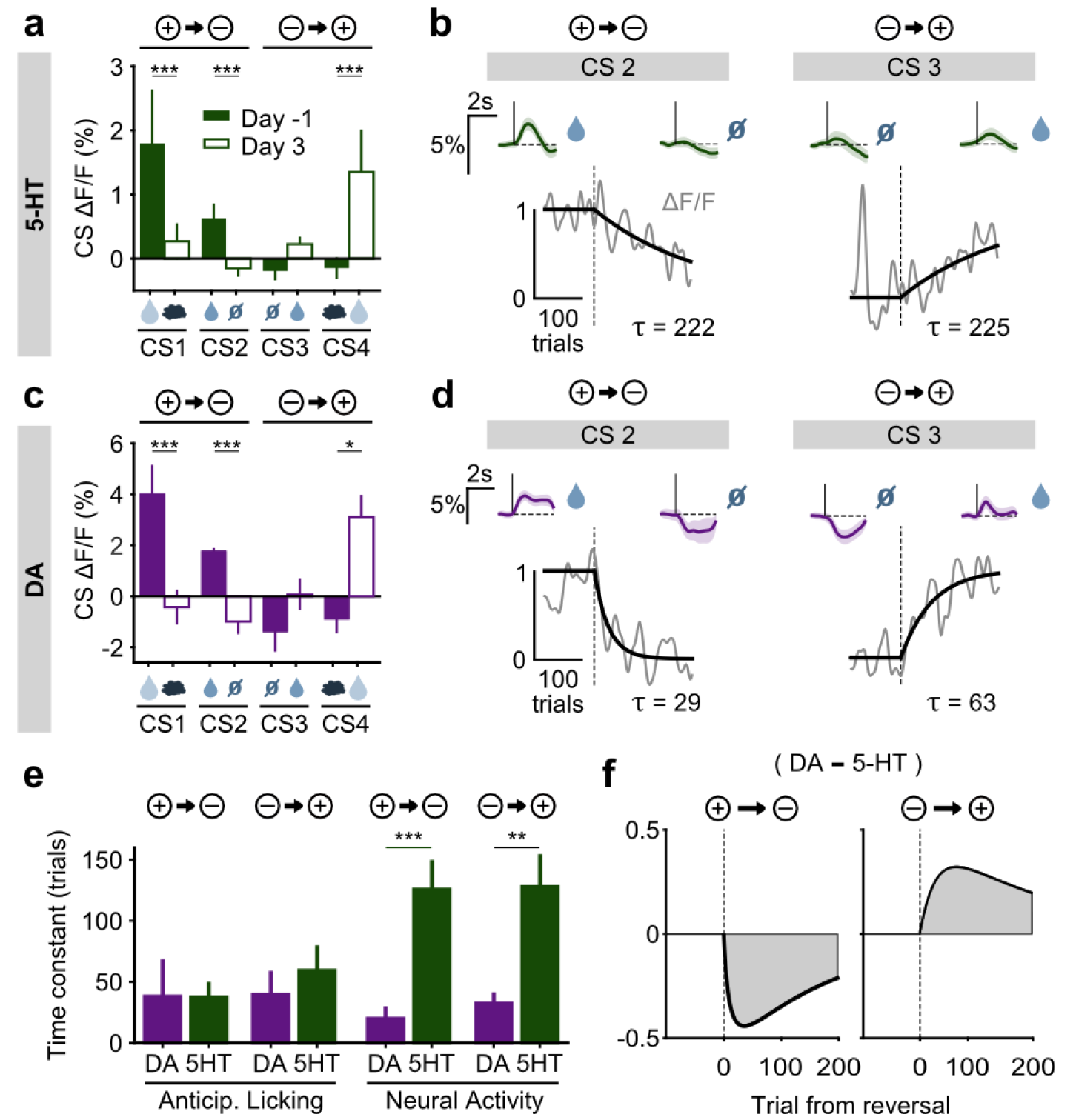
Distinct dynamics of CS responses in DRN 5-HT and midbrain DA neurons. **a**, Mean (± s.e.m.) response of 5-HT neurons across mice to the four CS’s before reversal (filled bars) and after adaptation to reversed contingencies (open bars) (n = 8 mice, 2-way ANOVA with factors day and mouse, main effect of day: large reward F_1.906_ = 17.35 *p* < 0.001, small reward F_1.902_ = 14.87 *p* < 0.001, neutral F_1.882_ = 0.13 *p* = 0.72, air puff F_1.914_ = 17.12 *p* < 0.001). **b**, Normalized exponential fits (black traces) to the mean amplitude of the CS responses (grey traces) across trials for CS 2 and CS 3 of an example SERT-Cre mouse. Insets show mean CS response (and 95% CI) on days -1 (left) and 3 (right). **c**, **d**, Same as **a**, **b** respectively for midbrain DA neurons (n = 3 mice, large reward F_1.294_ = 15.35 *p* < 0.001, small reward Fi_1.336_ = 71.72 *p* < 0.001, neutral F_1.282_ = 3.45, *p* = 0.06, air puff F_1.312_ = 6.56, *p* = 0.01). **e**, Mean time constants (± s.e.m.) of the exponential fits of CS responses obtained for TH-Cre and SERT-Cre mice during reversal learning (neural activity: unpaired *t*-tests *p* = 0.003 positive to negative reversal, *p* = 0.002 negative to positive reversal; no significance obtained for anticipatory licking). *f*, Difference in mean fitted amplitude of CS response between TH-Cre and SERT-Cre mice during negative reversal (left) and during positive reversal (right). \**p* < 0.05, \*\**p* < 0.01, \*\*\**p* < 0.001.

A potentially critical consequence of this difference in time constants is an asymmetry between positive and negative reversals (Fig. 4f). During a positive reversal, because the adaptation of 5-HT cue responses is much slower than that of DA cue responses, the net signal will be transiently biased towards the effects of DA (Fig. 4f, right). Conversely, during a negative reversal, because 5-HT cue responses persist longer than those of DA, the difference will be biased towards the effects of 5-HT (Fig. 4h, left). This suggests a novel mechanism by which 5-HT can contribute to preventing perseverative responding during negative reversals^4^.

Fibre photometry provides excellent genetic specificity and stable long-term recordings but does not allow the resolution of single neuron responses. It is therefore possible that differential activity patterns within specific subpopulations of DRN 5-HT neurons exist that could not be resolved by our methods. While this certainly merits further study, for several reasons we believe such heterogeneity would not substantially impact our conclusions. First, importantly, we established that the population from which we are recording contributes to reversal learning and it therefore is a relevant population. Second, activity patterns were consistent across mice (Extended Data Figs. 5,Fig. 8–Fig 10,Fig. 12,Fig. 13) despite inevitable small variations in infections and fibre placements across mice (Extended Data Fig. 2h), indicating that the findings are robust to the precise population monitored. Third, single unit recordings^22,29^ show that rewarding and aversive events activate the same individual DRN neurons and are thus not generated by distinct populations. Finally, because individual 5-HT neurons have broad projection fields^42^ and transmit primarily by volume conduction^43^, heterogeneity will tend to be averaged out through pooling by downstream targets.

The present findings support and extend a body of previous work implicating the 5-HT system in cognitive flexibility^3–8,10,13^. By recording the activity of a specific population of DRN 5-HT neurons involved in preventing perseverative responding during reversal learning, we found two different signals that provide insight into how the endogenous patterns of 5-HT activation contribute to this function. First, we found that 5-HT reports an unsigned prediction error, which is unsuitable to serve as reinforcement signal, but by signalling failures of prediction or control, is ideal to regulate learning, attention and effort^26,27,44^. Second, we found that cue-evoked 5-HT and DA signals are very similar, as reported previously^22^, but that 5- HT signals are much slower to adapt to changes in environmental contingency, an unappreciated difference with critical implications for their interaction during learning (Fig. 4f). Together, these findings suggest 5-HT is an affectively neutral modulator of plasticity and behaviour and suggest the possible need for a refinement in conventional conceptions of 5-HT in depression and other psychiatric disorders.

## Acknowledgements

We thank R. M. Costa and J. J. Paton Labs for TH-Cre mice; Susana Dias and Sérgio Casimiro for histology and immunohistochemistry assistance; Dario Sarra for running behavioural experiments for a few days. We also thank C. Poo, B. V. Atallah M. Murakami G. Agarwal and J. J. Paton for comments on a previous version of the manuscript, and all members of the Systems Neuroscience Lab and the Champalimaud Research for useful discussions and feedback during the development of this project. This work was supported by Fundação para a Ciência e Tecnologia (fellowship SFRH/BD/43072/2008 to S.M.), Human Frontier Science Program (fellowship LT000881/2011L to E.L.), European Research Council (Advanced Investigator Grants 250334 and 671251 to Z.F.M.), and Champalimaud Foundation (Z.F.M.).

## Author Contributions

S.M., E.L. and Z.F.M. designed the experiments and wrote the manuscript. S.M., G.P. D., and E.L. developed the fibre photometry with movement artefact correction technique. S.M. performed experiments. S.M. and E.L. performed data analysis.

## Supplemental Information

**Extended Data Figure 1.**
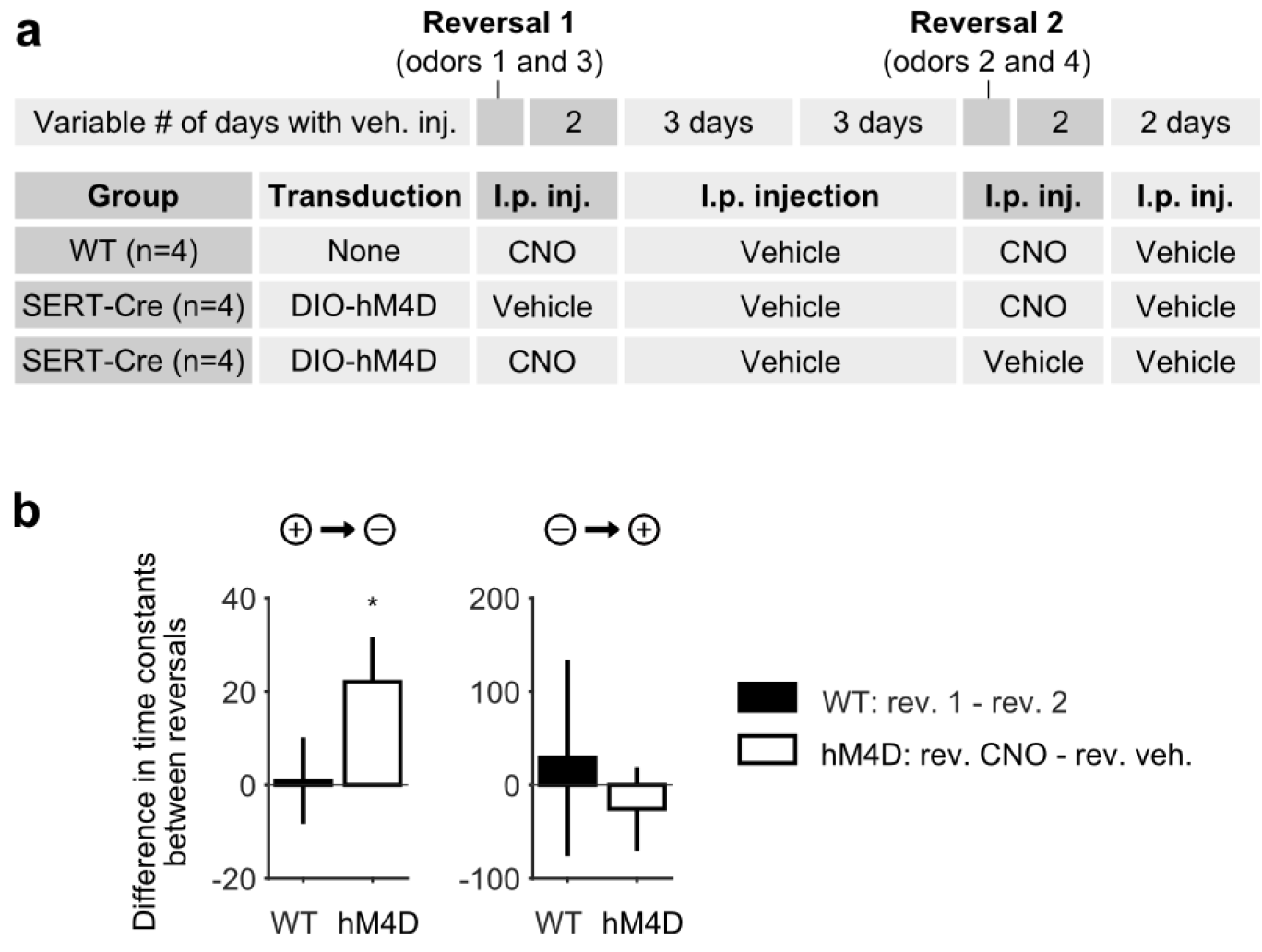
Anticipatory licking is more perseverative when DRN 5-HT neurons are inhibited. **a**, Detailed schematics of the manipulation experiment, including the viral transduction and i.p. injection received by experimental and control animals. b, In negative reversals (but not in positive reversals nor in WT controls receiving CNO in both reversals) adaptation in anticipatory licking takes longer when experimental animals receive CNO injection compared to when they receive vehicle (one-sample Student’s *t*-test). \**p* < 0.05.

**Extended Data Figure 2.**
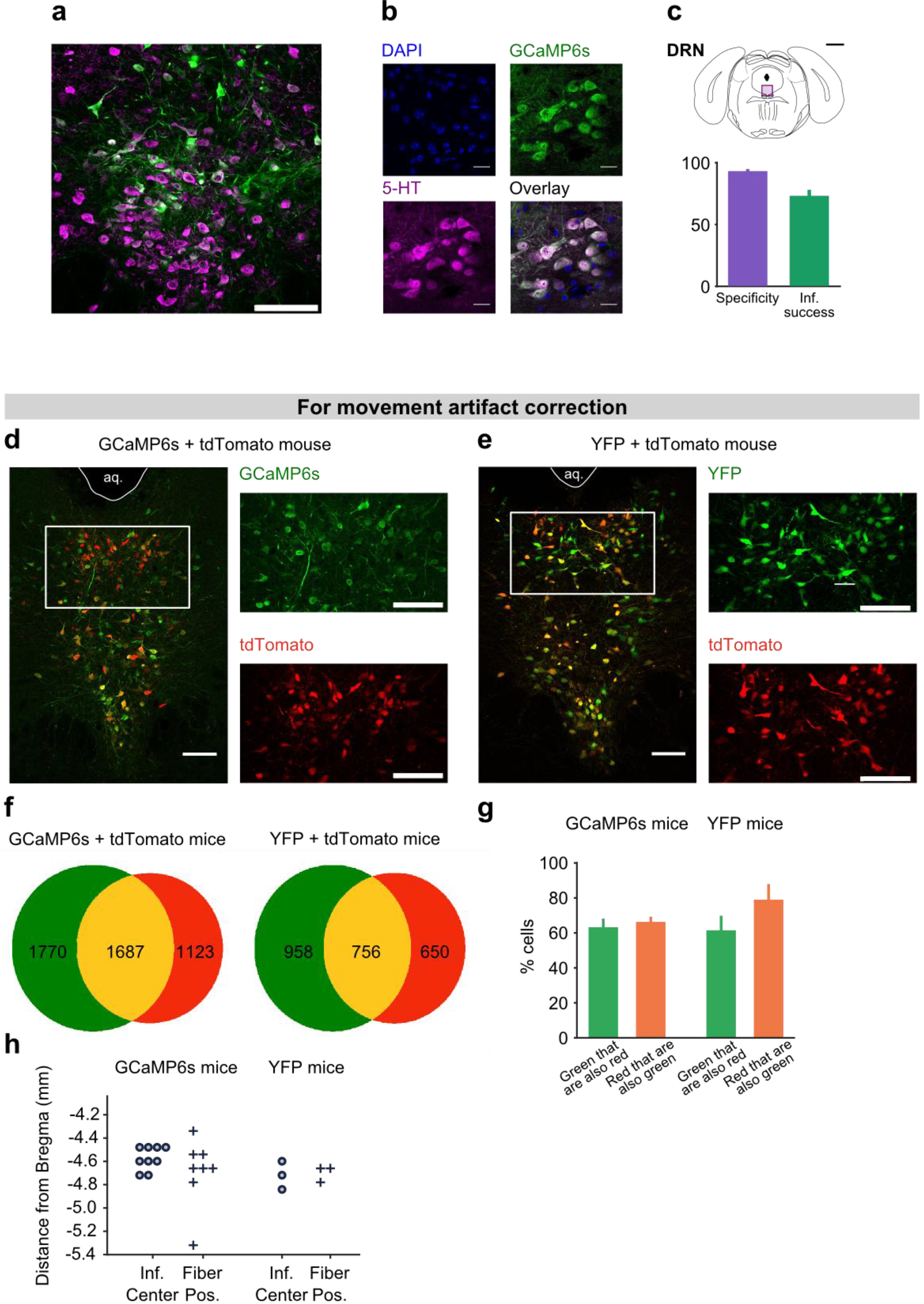
Expression of GCaMP6s and of tdTomato in DRN 5-HT neurons. **a**, Confocal picture of a coronal slice showing GCaMP6s expression in DRN 5-HT neurons of a SERT-Cre mouse (scale bar: 100 μm). **b**, Confocal pictures showing DAPI staining, GCaMP6s expression, 5-HT immunoreactivity and overlay (scale bar: 20 μm). **c**, Schematics of a coronal slice showing the DRN and the area in **a** signaled by a violet rectangle (top, black scale bar: 1 mm). Quantification of specific expression of GCaMP6s in 5-HT neurons (bottom, specificity: 93.2 ± 0.3 %, infection success: 73.3 ± 3.5 %, mean ± s.e.m, n = 4 mice). **d**, Expression of GCaMP6s and tdTomato in DRN 5-HT neurons (scale bars: 100 μm aq. - aqueduct). **e**, Expression of YFP and tdTomato in DRN 5-HT neurons (scale bars: 100 μm). **f**, Total number of cells expressing green fluorophore (GCaMP6s or YFP), tdTomato, or both in SERT-Cre experimental and control mice (counted from 3 sections at the center of infection for each mouse, n = 6 GCaMP6s mice, n = 3 YFP mice). **g**, Percent of green and red cells that express both fluorophores in GCaMP6s and YFP infected mice (n = 6 GCaMP6s mice, n = 3 YFP mice, no statistical difference was obtained between the percentage of green cells that co-express tdTomato in GCaMP vs YFP mice, nor between the percentage of red cells that co-express a green fluorophore in GCaMP vs YFP mice, Mann-Whitney U test n.s. for *p* < 0.05). **h**, Antero-posterior location of the center of infection (circles) and of the fibre tip location (crosses) in the SERT-Cre mice used in behavioral experiments with good histology (n = 9 for GCaMP6s infection center, n = 8 for GCaMP6s fiber placement, n = 3 for YFP control mice).

**Extended Data Figure 3.**
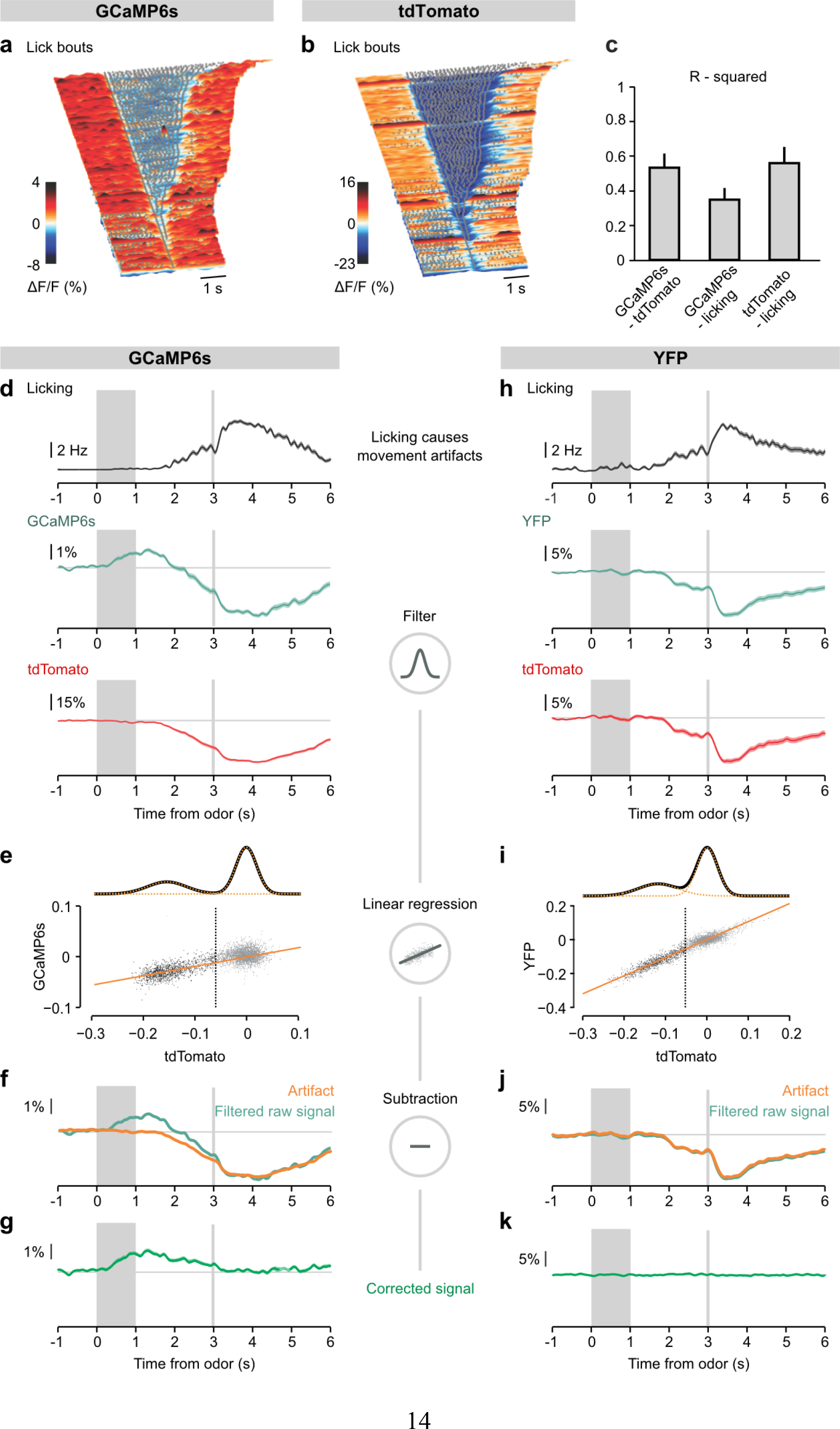
Linear regression approach to eliminate movement artefacts from neuronal photometric data. **a, b**, Surface plots showing raw GCaMP6s (a) and tdTomato (b) fluorescence signals aligned on the onsets of licking bouts during an example session of a SERT-Cre mouse. Gray dots represent single licks. Bouts were defined as sequences of licks separated by no more than 315 ms. c, Bar plot showing R-squared values comparing GCaMP6s, tdTomato and licking signals calculated during 3 imaging sessions and averaged across all SERT-Cre mice (n = 9). **d**, **h** Mean licking (top), mean filtered green fluorescence of GCaMP6s (d) or YFP (h), and mean filtered tdTomato fluorescence in large water reward trials in an example session of a SERT-Cre mouse transduced with GCaMP6s and tdTomato and of a SERT-Cre example mouse transduced with YFP and tdTomato. Data in (d) belongs to the same session represented in (a,b). **e**, **i**, Scatter plot and linear regression between tdTomato and GCaMP6s (e) or YFP (i) signals for the same sessions as before. tdTomato signals were often bimodal and well fit as a sum of 2 Gaussians (top; orange dotted curves, individual Gaussians; black curve, their sum). In order to avoid noise fitting, only data from the Gaussian not centered at 0 were included in the linear regression calculation (black dots to the left of the vertical line). Orange lines indicate regression curves. **f**, **j**, Mean filtered raw GCaMP6s (f) or YFP (j) signals and corresponding artefact predictions (calculated using linear regression as shown in *e* and *i*). **g**, **k**, Corrected fluorescence signal obtained after the subtraction of the filtered raw signal by the artefact signal presented in (f) and (j). While a calcium signal is still visible in the mouse infected with GCaMP6s, no signal is observed in the YFP control.

**Extended Data Figure 4.**
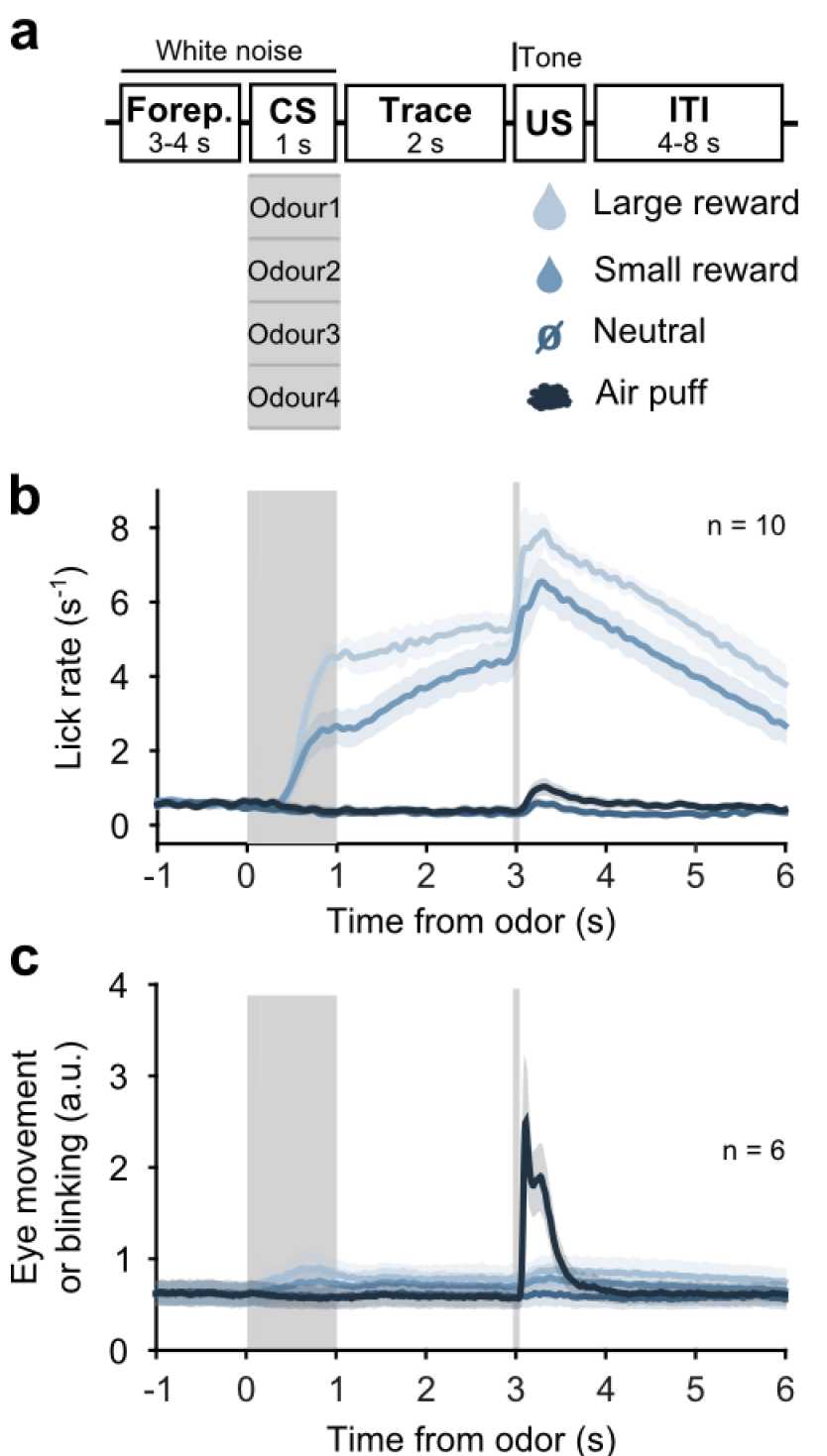
Behavior of head-fixed mice trained in a classical conditioning task. **a**, Schematics of the trial structure in the classical conditioning task (before reversal) with four different outcomes. In each trial one of four odours was randomly selected and presented during 1 s after a variable foreperiod (Forep). The associated outcome was delivered after a 2 s trace period together with a tone (same tone for all trial types). Mice were presented with 140 to 346 interleaved trials (mean ± SD: 223 ± 30) per session(day). **b**, Mean licking rate of SERT-Cre mice infected with GCaMP6s and tdTomato used in this task (n = 10) along the duration of each trial type. For each mouse, three sessions of the classical conditioning task where initial associations had already been learned were averaged. Colors code trial type according to (1a), shaded areas represent s.e.m. **c**, Mean eye movement across SERT-Cre mice infected with GCaMP6s and tdTomato (n = 6) along the duration of each trial type.

**Extended Data Figure 5.**
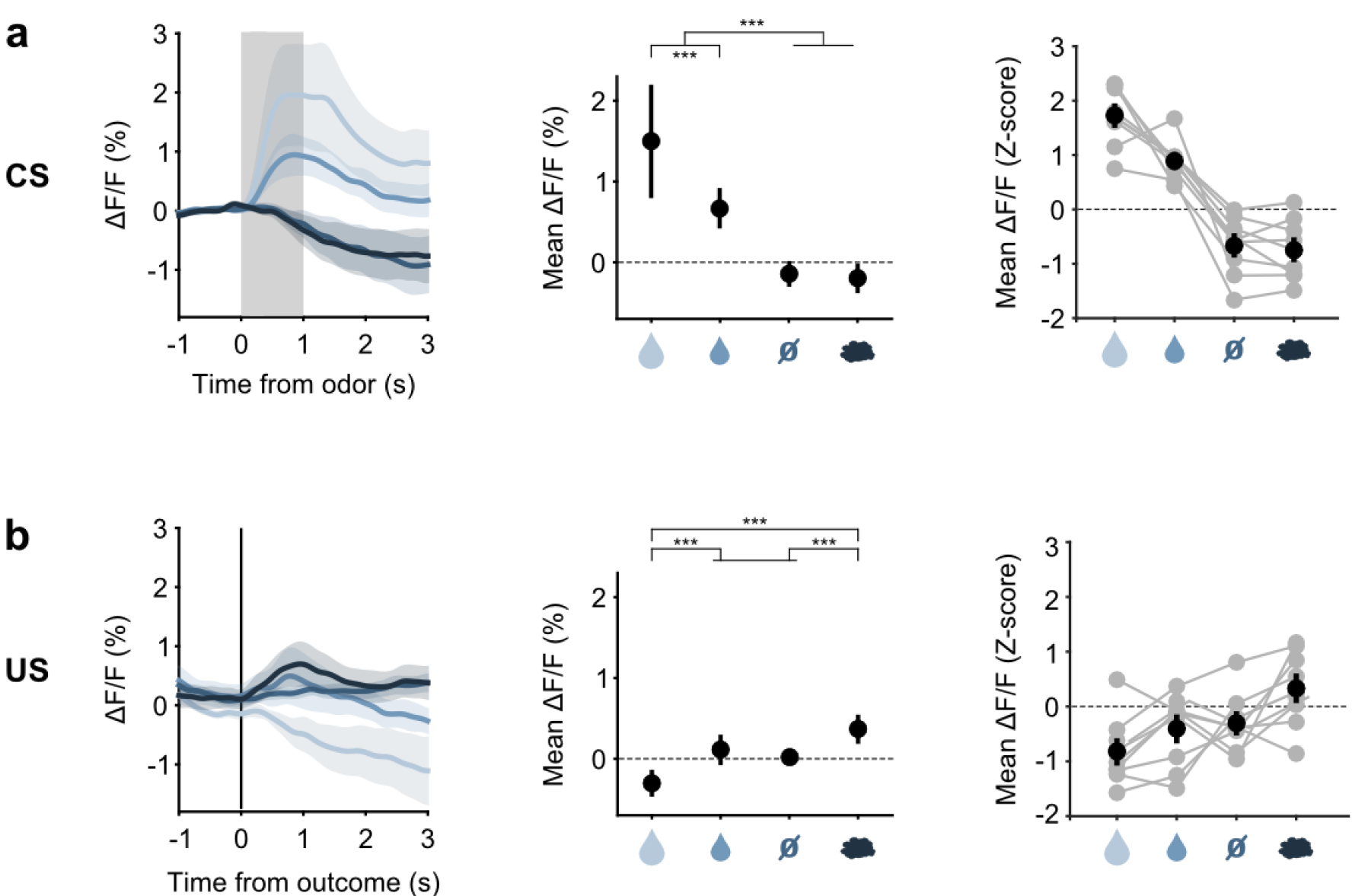
Responses of DRN 5-HT neurons to odour cues and to predicted outcomes. **a**, Left: Mean (± s.e.m) CS responses (corrected fluorescence) of all SERT-Cre mice expressing GCaMP6s (n = 9) to the four odours (CS’s) after having learned the CS-US associations; Middle: quantification of the mean (± s.e.m) CS response amplitude (in a 1.5 s period from odour onset) across mice (3-way ANOVA with factors day, mouse and trial type, main effect of trial type F_3,6365_ = 362.78, *p* < 0.001, only statistically significant *pos hoc* multiple comparisons for trial type are shown: vertical ticks signal events being compared, horizontal bars without vertical ticks below them are used for grouping non-statistically different events); Right: Z-scores of mean response amplitude for individual mice (grey dots, n = 9 mice) and their mean (± s.e.m, black dots). **b**, Same as (a) but for US responses of SERT-Cre mice (F_3,6365_ = 55.62, *p* < 0.001). The predicted large reward US responses of 5-HT neurons have a slightly negative value because of the response that the corresponding CS triggered before and which takes longer to go back to baseline than the 2 s trace period. \**p* < 0.05, \*\**p* < 0.01, \*\*\**p* < 0.001.

**Extended Data Figure 6.**
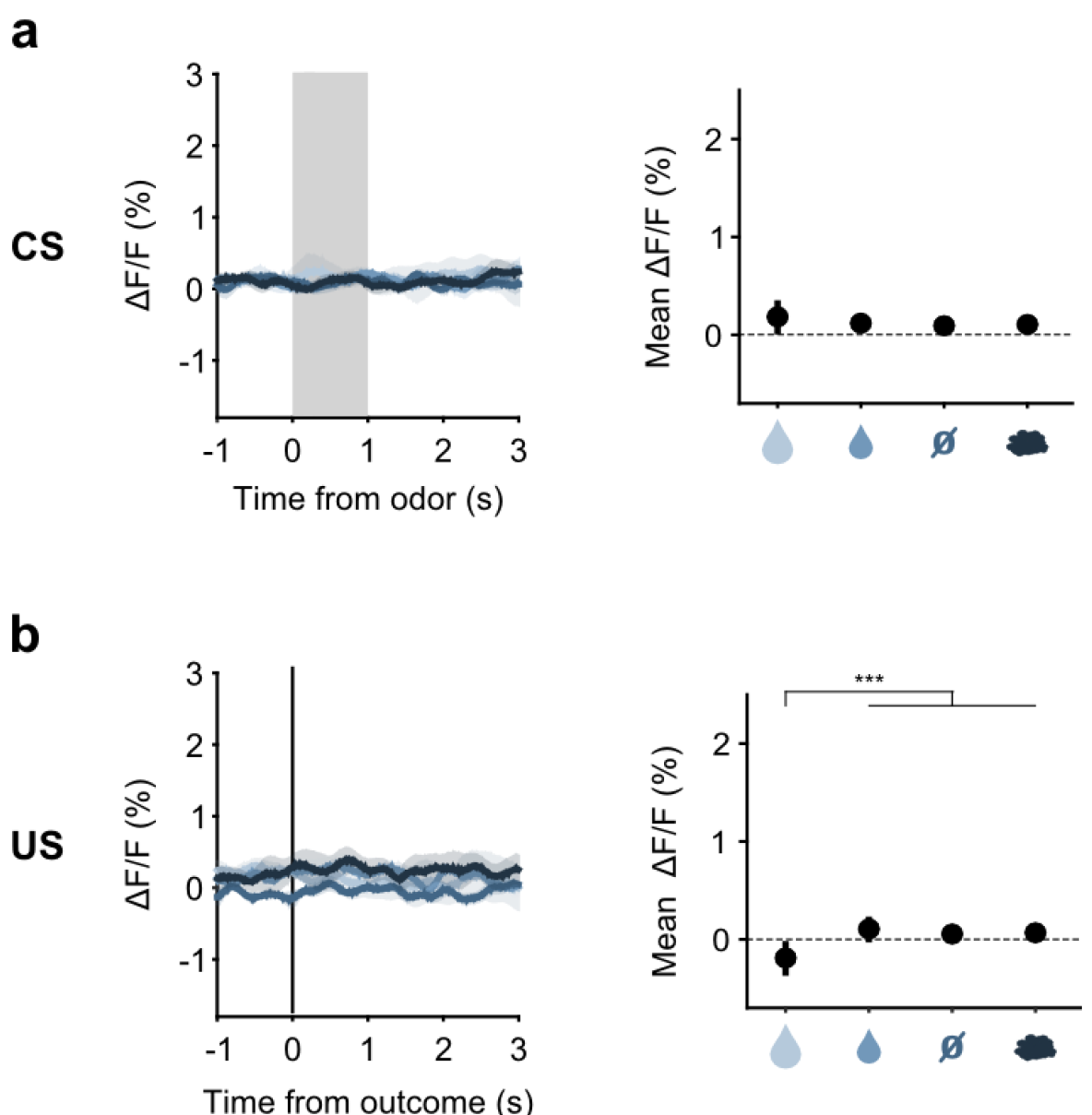
YFP controls for DRN 5-HT neurons fluorescence changes to odour cues and to predicted outcomes. **a**, Left: Mean (± s.e.m.) CS responses (corrected fluorescence) of all SERT-Cre mice infected with YFP instead of GCaMP6s (n = 4) to the four odours after having learned the CS-US associations; Right: quantification of the mean (± s.e.m.) CS response amplitude (in a 1.5 s period from odour onset) across mice (3-way ANOVA with factors day, mouse and trial type, main effect of trial type F_3,2631_ = 1.4, *p* = 0.2412). **b**, Same as (a) but for US responses of YFP-expressing SERT-Cre mice (F_3,2631_ = 11.95, *p* < 0.001).\**p* < 0.05, \*\**p* < 0.01, \*\*\**p* < 0.001.

**Extended Data Figure 7.**
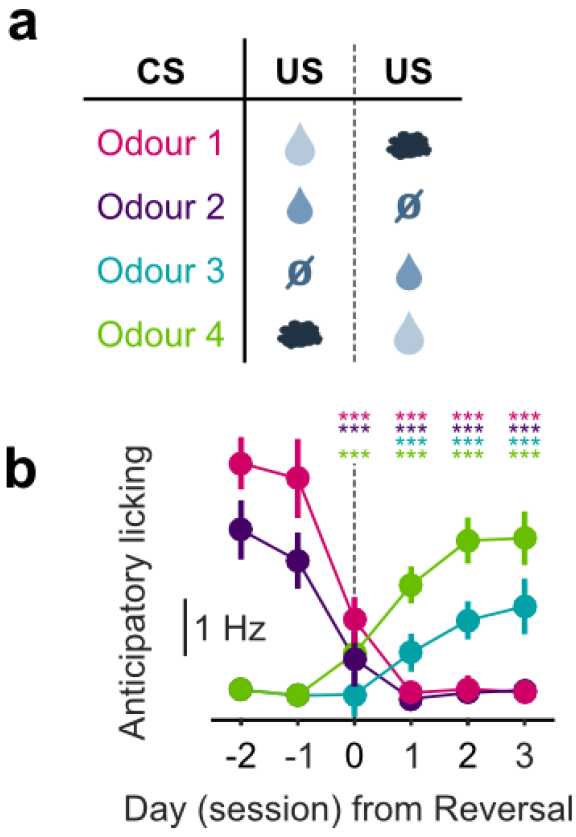
Mice adjust their anticipatory licking after reversal. **a**, Reversal of CS-US associations. **b**, Anticipatory licking (mean of 500-2800 ms after odour onset) across mice for sessions around reversal showing that the licking rate triggered by the presentation of each odour is adjusted after reversal (n = 8, 2-way ANOVA with factors day (days -2 and -1 are considered together) and mouse, main effect of day: F_4,2597_ = 722.14 *p* < 0.001 for odour 1, F_4,2554_ = 355.53 *p* < 0.001 for odour 2, F_4,2513_ = 104.93 *p* < 0.001 for odour 3, F_4,2559_ = 381.55 *p* < 0.001 for odour 4). Colors follow odour identity as in (a). \*\*\**p* < 0.001.

**Extended Data Figure 8.**
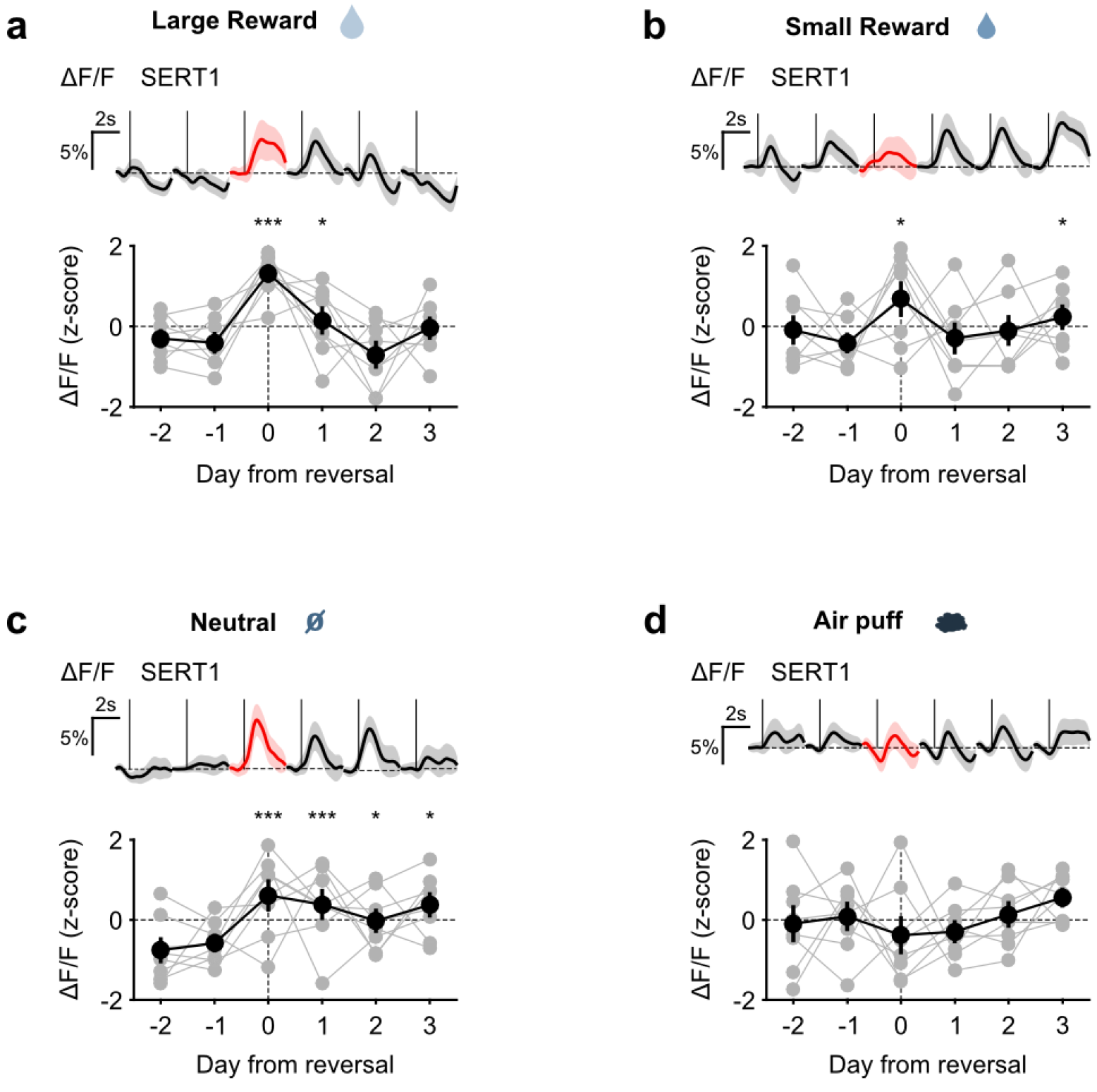
US responses of DRN 5-HT neurons during reversal. **a**, Top: Mean large reward US responses of an example mouse (SERT1) over days around reversal (shaded areas represent 95% CI); Bottom: change in mean large reward response amplitude (z-scored across days): grey dots represent individual mice (n = 8), black dots average (± s.e.m.) of mice (2-way ANOVA with factors day and mouse, main effect of day F_4,2592_ = 31.47 *p* < 0.001 for large reward; multiple comparisons with 2 days before reversal using Scheffé’s method are signaled in the figure). **b, c, d**, Same as (a) for small reward US (F_4,2532_ = 5.98, *p* < 0.001), neutral US (F_4,2535_ = 10.71, *p* < 0.001), and air puff US (F_4,2564_ = 2.55, *p* = 0.037) respectively. \**p* < 0.05, \*\**p* < 0.01, \*\*\**p* < 0.001.

**Extended Data Figure 9.**
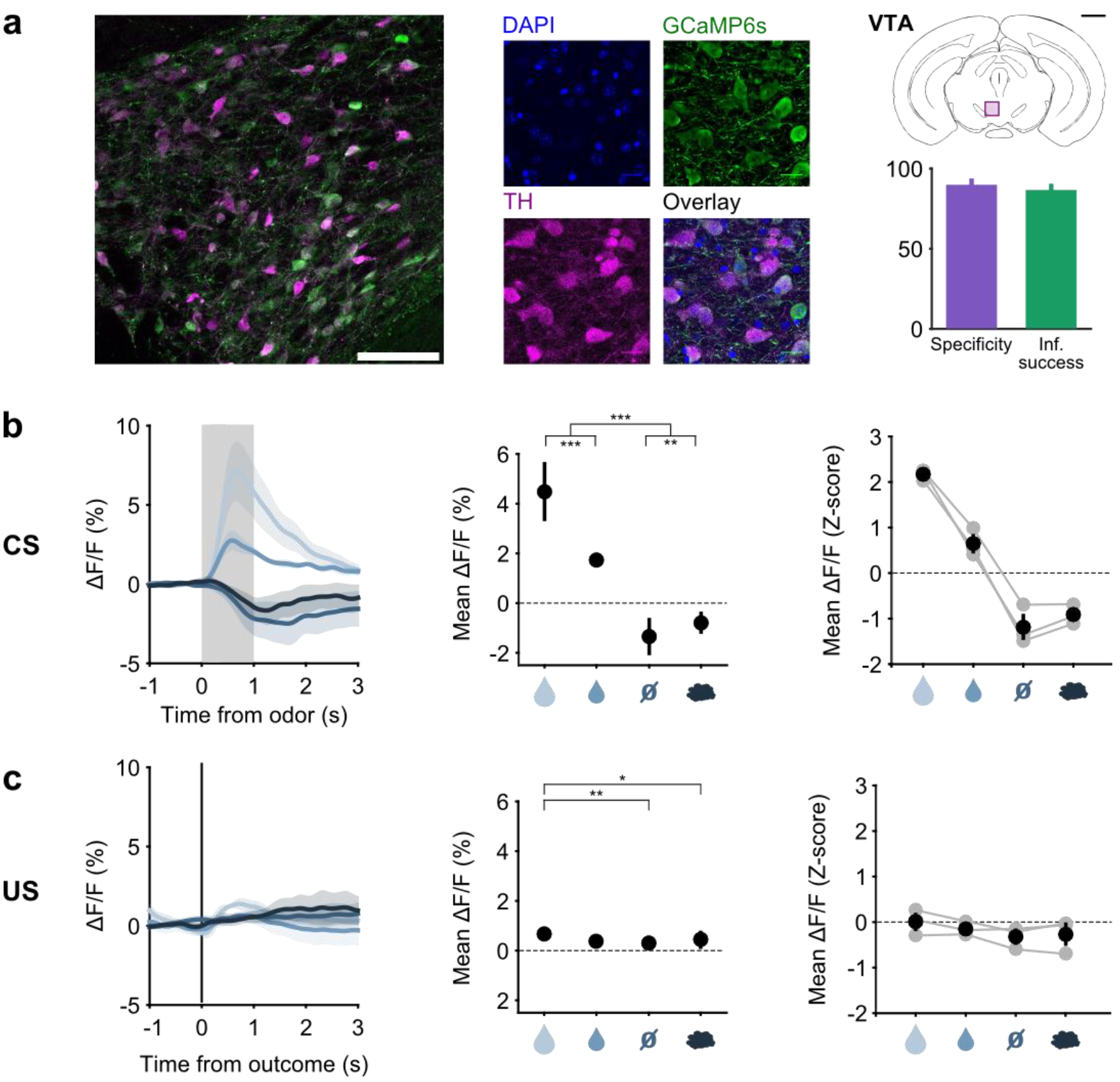
Responses of midbrain DA neurons before reversal. **a**, Confocal picture of a coronal slice showing GCaMP6s expression in VTA DA neurons of a TH-Cre mouse (left, scale bar: 100 μm); close up showing DAPI staining, GCaMP6s expression, TH immunoreactivity and overlay (middle, scale bar: 20 μm); schematics of a coronal slice showing the VTA and the area on the left signaled by a violet rectangle (scale bar: 1 mm) with quantification of specific expression of GCaMP6s in DA neurons in the infection center (right, specificity: 89.9 ± 2.5 %, infection success: 86.7 ± 2.5 %, mean ± s.e.m, n = 4 mice). **b**, Left: Mean (± s.e.m) CS responses (corrected fluorescence) of all TH-Cre mice expressing GCaMP6s (n = 3) to the four odours (CS’s) after having learned the CS-US associations; Middle: quantification of the mean (± s.e.m) CS response amplitude (in a 1.5 s period from odour onset) across mice (3-way ANOVA with factors day, mouse and trial type, main effect of trial type F_3,1832_ = 671.64 *p* < 0.001, only statistically significant *pos hoc* multiple comparisons for trial type are shown: vertical ticks signal events being compared, horizontal bars without vertical ticks below them are used for grouping non-statistically different events); Right: Z-scores of mean response amplitude for individual mice (grey dots, n = 3 mice) and their mean (± s.e.m, black dots). **b**, Same as (b) but for US responses of TH-Cre mice (F_3,1832_ = 5.52, *p* < 0.001). \**p* < 0.05, \*\**p* < 0.01, \*\*\**p* < 0.001.

**Extended Data Figure 10.**
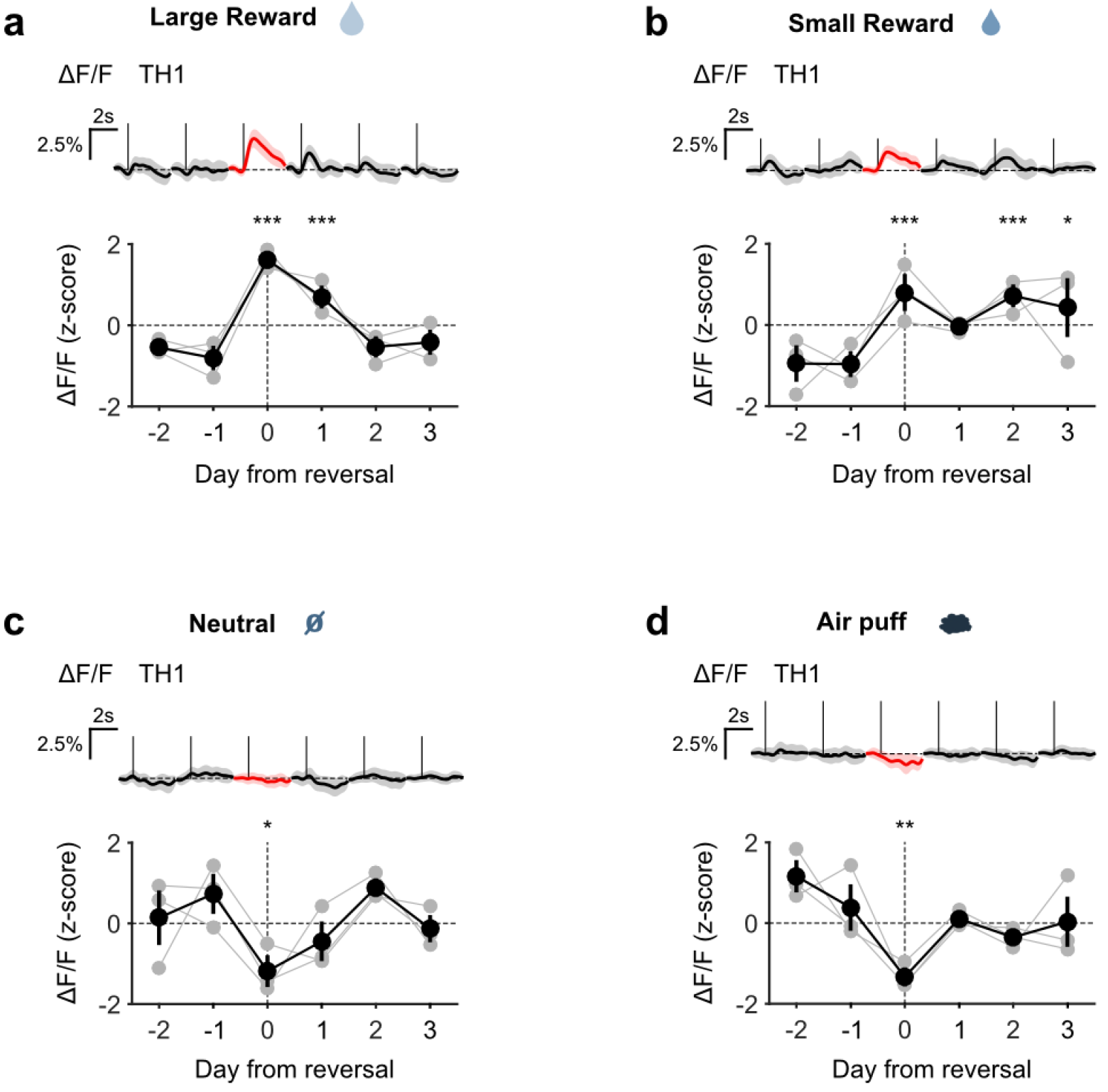
US responses of midbrain DA neurons during reversal. Top: Mean large reward US responses of an example mouse (TH1) over days around reversal (shaded areas represent 95% CI); Bottom: change in mean large reward response amplitude (z-scored across days): grey dots represent individual mice (n = 3), black dots average (± s.e.m.) of mice (2- way ANOVA with factors mouse and day, main effect of day F_4,853_ = 32.46, *p* < 0.001; multiple comparisons with 2 days before reversal using Scheffé’s method are signaled in the figure). **b, c, d**, Same as (a) for small reward US (F_4,911_ = 8.52, *p* < 0.001), neutral US (F_4,843_ = 4.54, *p* = 0.001), and air puff US (F_4,881_ = 5.21, *p* < 0.001) respectively.\**p* < 0.05, \*\**p* < 0.01, \*\*\**p* < 0.001.

**Extended Data Figure 11.**
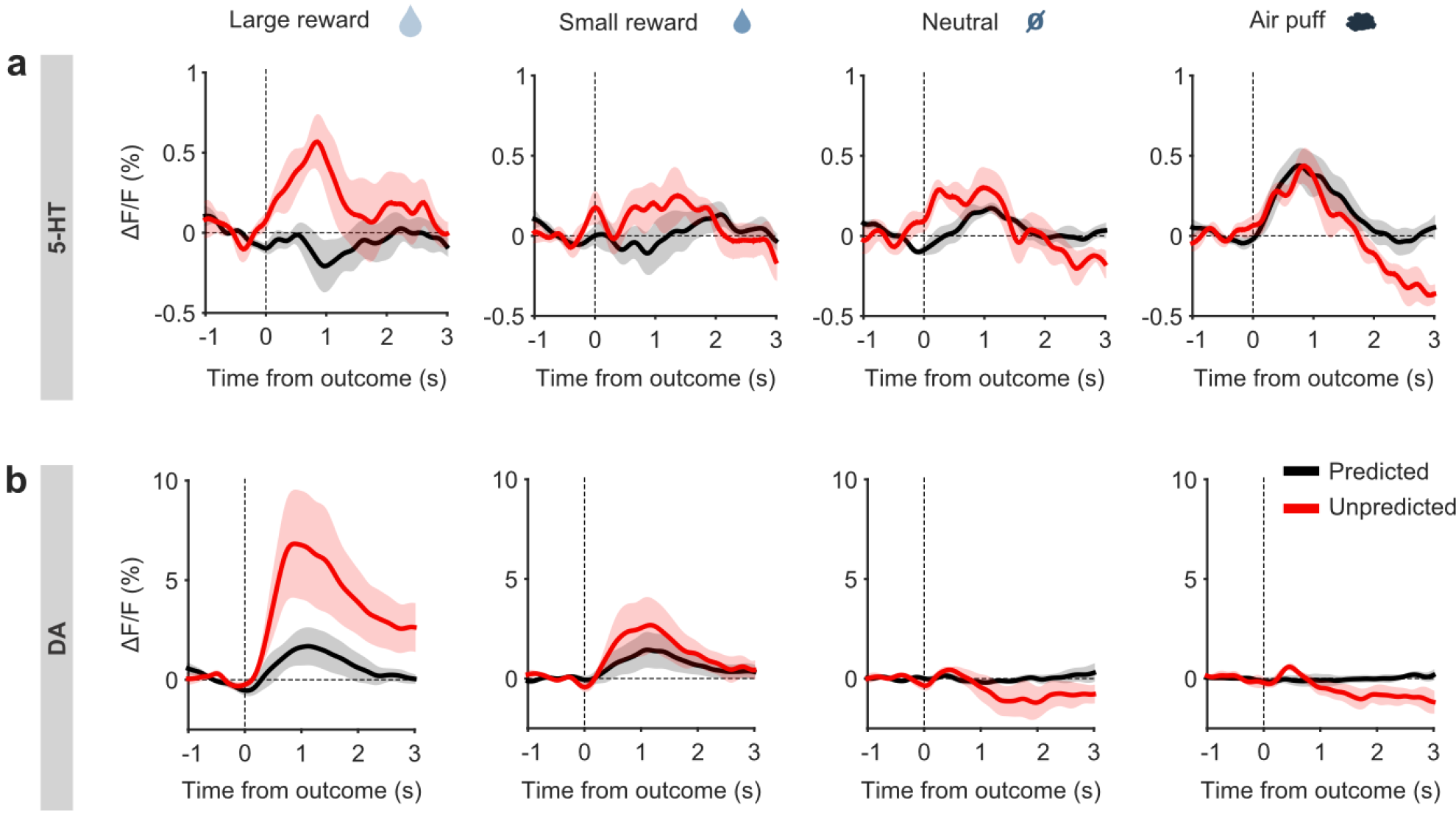
Responses of DA and 5-HT neurons to surprising outcomes. **a**, Mean US responses of SERT-Cre mice (n = 4) to predicted (black) and unpredicted (red) outcomes (shaded areas represent s.e.m.). b, Same as (a) for TH-Cre mice.

**Extended Data Figure 12.**
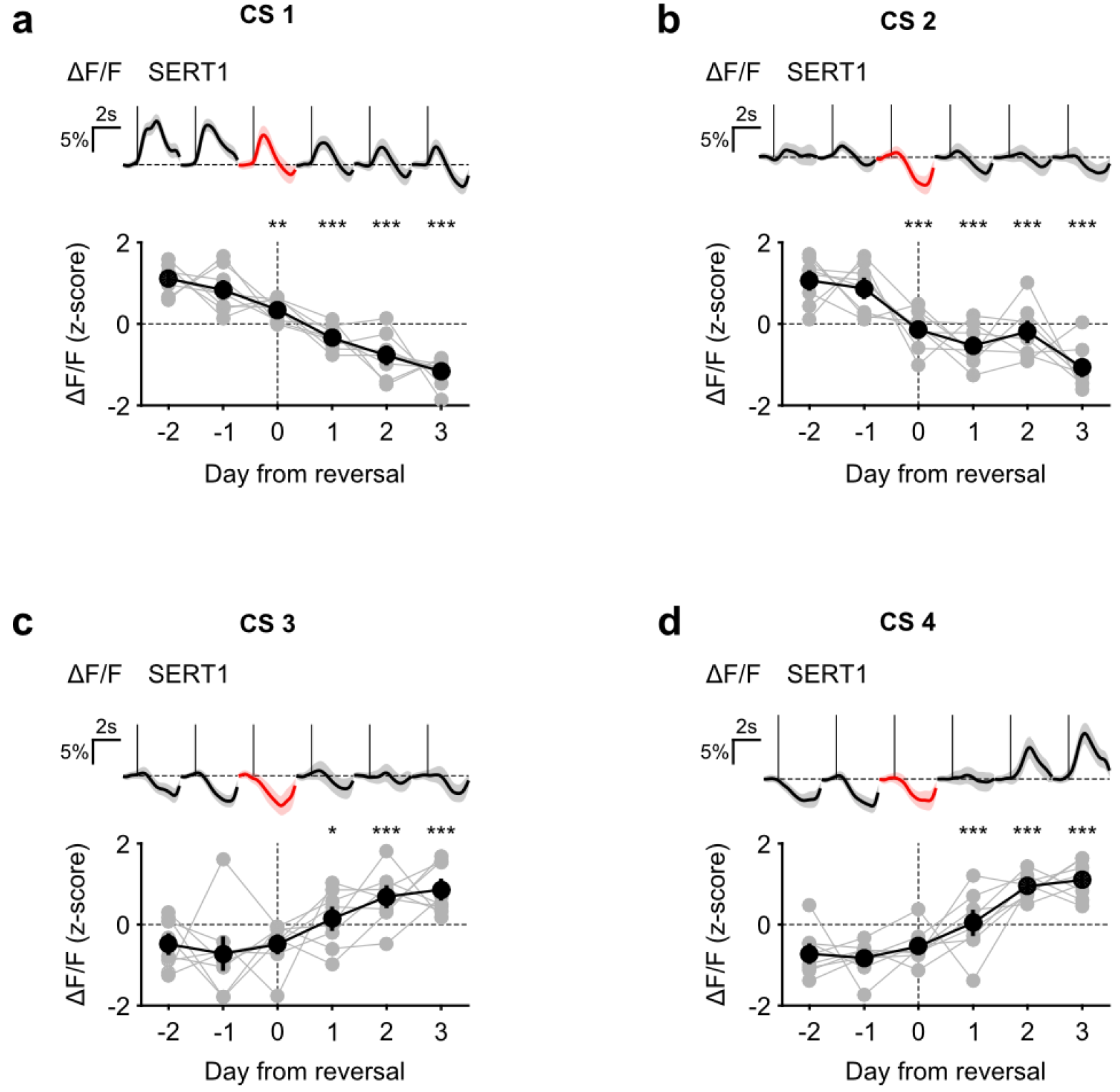
CS responses of DRN 5-HT neurons during reversal. Top: Mean responses of an example mouse (SERT1) to odour 1 over days around reversal (shaded areas represent 95% CI); Bottom: change in mean response amplitude to odour 1 (z-scored across days): grey dots represent individual mice (n = 8), black dots average (± s.e.m.) of mice (2- way ANOVA with factors day and mouse, main effect of day F_4,2597_ = 72.1, *p* < 0.001; multiple comparisons with 2 days before reversal using Scheffé’s method are signaled in the figure). **b, c, d**, Same as (a) for small reward US (F_4,2554_ = 33.37, *p* < 0.001), neutral US (F_4,2513_ = 10.88, *p* < 0.001), and air puff US (F_4,2559_ = 73.36, *p* < 0.001) respectively. \**p* < 0.05, \*\**p* < 0.01, \*\*\**p* < 0.001.

**Extended Data Figure 13.**
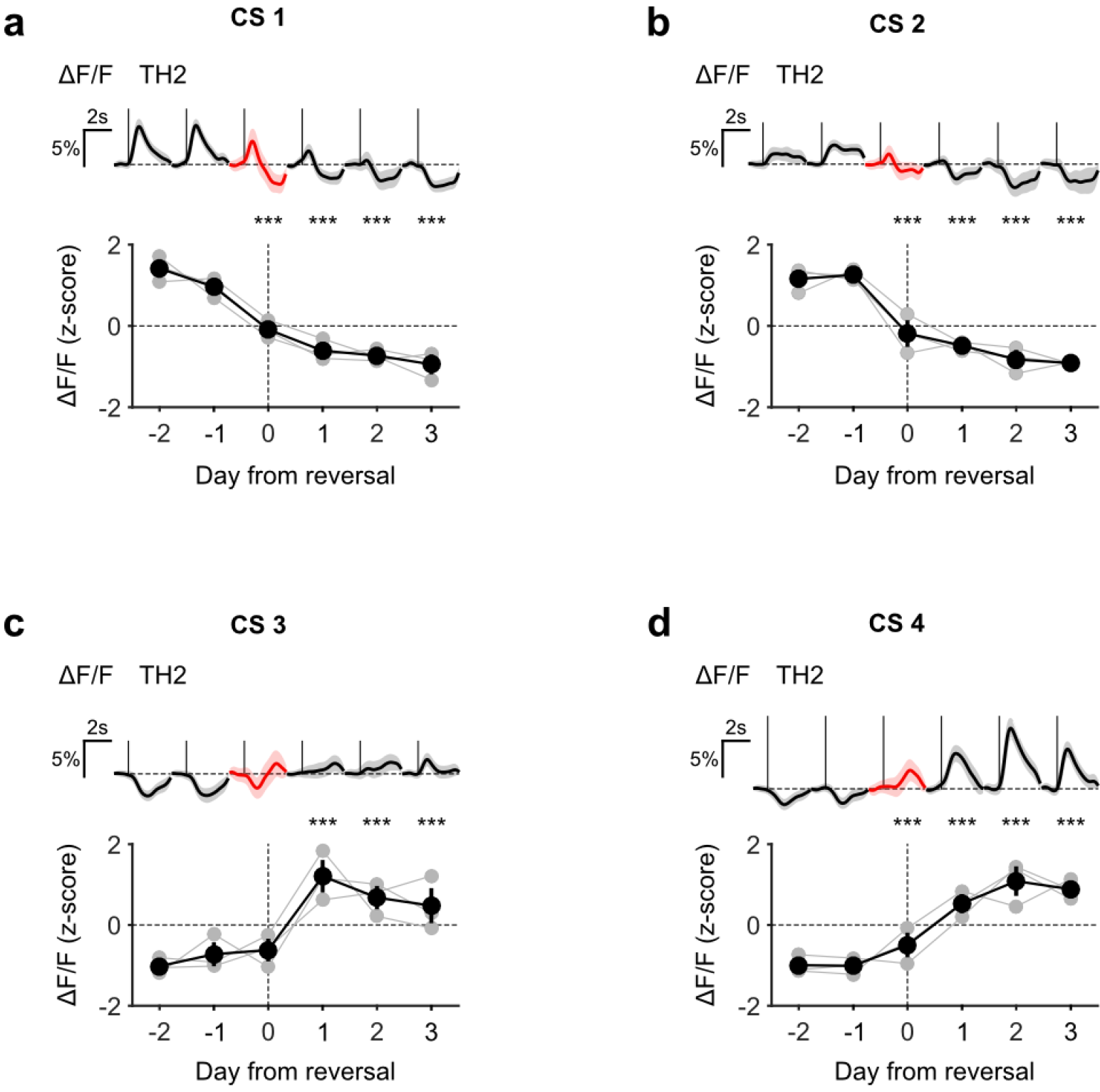
CS responses of midbrain DA neurons during reversal. Top: Mean large reward US responses of an example mouse (TH2) over days around reversal (shaded areas represent 95% CI); Bottom: change in mean large reward response amplitude (z-scored across days): grey dots represent individual mice (n = 3), black dots average (± s.e.m.) of mice (2-way ANOVA with factors day and mouse, main effect of day F_4,882_ = 223.44, *p* < 0.001; multiple comparisons with 2 days before reversal using Scheffé’s method are signaled in the figure). **b, c, d**, Same as (a) for small reward US (F_4,856_ = 84.97, *p* < 0.001), neutral US (F_4,898_ = 29.86, *p* < 0.001), and air puff US (F_4,852_ = 151.95, *p* < 0.001) respectively. \**p* < 0.05, \*\**p* < 0.01, \*\*\**p* < 0.001.

## Methods

**Animals** - All procedures were reviewed and performed in accordance with the Instituto Gulbenkian de Ciência and the Champalimaud Centre for the Unknown Ethics Committee guidelines, and approved by the Portuguese Veterinary General Board (Direco Geral de Veterinaria, approval ID 018831). Thirty four C57BL/6 male mice between 2 and 9 months of age were used in this study. Mice resulted from the backcrossing of BAC transgenic mice into Black C57BL for at least six generations, which express the Cre recombinase under the control of specific promoters. Twenty six mice expressed Cre under the serotonin transporter gene (Tg(Slc6a4-cre)ET33Gsat/Mmucd) from GENSAT^33^, and 4 mice under the tyrosine hydroxylase gene, 2 mice (Tg(Th-cre)FI12Gsat/Mmucd) from GENSAT^33^, and 2 mice (B6.Cg-Tg(Th-Cre)1Tmd/J) from The Jackson Laboratory^45^. Animals (25 - 45 g) were group-housed prior to surgery and individually housed post-surgery and kept under a normal 12 hour light/dark cycle. All experiments were performed in the light phase. Mice had free access to food. After training initiation, mice used in behavioural experiments had water availability restricted to the behavioural sessions.

**Stereotaxic viral injections and fibre implantation** - Mice were deeply anaesthetized with isoflurane mixed with O_2_ (4% for induction and 0.5-1% for maintenance) and placed in a stereotaxic apparatus (David Kopf Instruments). Butorphanol (0.4 mg/kg) was injected subcutaneously for analgesia and Lidocaine (2%) was injected subcutaneously before incising the scalp and exposing the skull. For SERT-Cre mice a craniotomy was drilled over lobule 4/5 of the cerebellum and a pipette filled with a viral solution was lowered to the DRN (bregma −4.55 anterioposterior (AP), −2.85 dorsoventral (DV)) with a 32° angle toward the back of the animal. For the 2 TH-Cre mice from The Jackson Laboratory the pipette was targeted to the VTA (bregma −3.3 AP, 0.35 mediolateral (ML), −4.2 DV) with a 10° lateral angle, and for the 2 TH-Cre mice from GENSAT we targeted the SNc (bregma - 3.15 AP, 1.4 ML, −4.2 DV). Although the TH-Cre lines have been characterized as less specific than other DA-specific lines^46^, we targeted our fibres to areas where this specificity problem is reduced^46^ and that are known to contain the classical DA neurons that show RPE activity and are involved in reward processing^38-40,47,48^.

Viral solution was injected using a Picospritzer II (Parker Hannifin) at a rate of approximately 38 nl per minute. The expression of hM4D and of all fluorophores was Cre-dependent and all viruses were obtained from University of Pennsylvania (with 10^12^ or 10^13^ GC/mL titers). For hM4D experiments 1 μl AAV2/5 - Syn.DIO.hM4D.mCherry was injected in the DRN of 8 SERT-Cre mice. For analysis of GCaMP6s specific expression in 5-HT neurons 4 SERT-Cre mice were transduced in the DRN with 1 μl of viral stock solution of AAV2/1 - Syn.Flex.GCaMP6s.WPRE.SV40. For behavioural experiments in control mice (4 SERT-Cre mice) 1.5 μl of a mixture of equal volumes of AAV2/1.EF1a.DIO.eYFP.WPRE.hGH and of AAV2/1.CAG.FLEX.tdTomato.WPRE.bGH was used. The remaining mice were injected with a mixture of equal volumes of AAV2/(1 or 9).Syn.Flex.GCaMP6s.WPRE.SV40 and of AAV2/1.CAG.FLEX.tdTomato.WPRE.bGH: 1.5 μl was injected in 10 SERT-Cre mice (distributed around 6 points around the target coordinates) and 0.75 μl of 10 times diluted mixture was injected in 4 TH-Cre mice (distributed around 4 points around the target).

For photometry experiments, optical fibre implantations were done after infection and a head plate for head fixation was placed above Bregma: the skull was cleaned and covered with a layer of Super Bond C&B (Morita). An optical fibre (300 μm, 0.22 NA) housed inside a connectorized implant (M3, Doric Lenses) was inserted in the brain, with the fibre tip positioned at the target for SERT-Cre mice and 200 μm above the infection target for TH-Cre mice. The implants were secured with dental acrylic (Pi-Ku-Plast HP 36, Bredent).

**Behavioural training and testing protocol** - Mice were water deprived in their homecage on the day of surgery or up to 5 days before it. During water deprivation mice weight was maintained above 80% of their original values. Following infection and implantation surgery, mice were habituated to the head-fixed setup by receiving water every 4s (6 μl drops) for 3 days, after which training in the odour-guided task started. A mouse poke (007120.0002, Island Motion Corporation) was adapted and used as a lickometer; sounds signaling the beginning of the trial and the outcomes were amplified (PCA1, PYLE Audio Inc.) and presented through speakers (Neo3 PDRW W/BC, Bohlender-Graebener), water valves (LHDA1233115H, The Lee Company) were calibrated and a custom made olfactometer designed by Z.F.M. (Island Motion) was used for odour delivery. The behavioural control system (Bcontrol) was developed by Carlos Brody (Princeton University) in collaboration with Calin Culianu, Tony Zador (Cold Spring Harbor Laboratory) and Z.F.M. Odours were diluted in mineral oil (Sigma-Aldrich) at 1:10 and 25 μl of each diluted odour was placed inside a syringe filter (2.7 μm pore size, 6823-1327, GE Healthcare) to be used in two sessions (~100 trials for each odour). Odourized air was delivered at 1000 ml.min^−1^. Odours used were carvone (R)-(−), 2-octanol (S)-(+), amyl acetate and cuminaldehyde. For the behavioural task used in the hM4D experiment these odours were associated with reward, reward, nothing and nothing respectively. For the behavioural task used in the GCaMP6s experiment, they were associated with large reward (4 μl water drop), small reward (2 μl water drop), neutral (no outcome) and punishment (air puff to the eye) before reversal, and with punishment, neutral, small reward and big reward after reversal of the cue-outcome associations respectively. In each trial, white noise was played to signal the beginning of the trial and to mask odour valve sounds. A randomly selected odour was presented during 1s. Following a 2s trace period the corresponding outcome was available. Mice did one session per day. For hM4D experiments odours were introduced in pairs. For photometry experiments training started by presenting only with the large and small reward trials to mice, followed by the introduction of the neutral type of trial in the next session and finally the punishment trial in the following one. Punishment trials were presented gradually until all 4 types of trials had the same probability of occurrence and each session consisted of 140 - 346 trials (minimum to maximum, 223 ± 30 mean ± SD). Time to odour (foreperiod), trace period and ITI were also gradually increased during training until mice could do the task with their final values: foreperiod was 3 to 4 s, taken from a uniform distribution, trace was fixed at 2 s, ITI was 4 to 8 s taken from a uniform distribution.

For the hM4D experiments mice received a daily injection of vehicle (saline 0.9% and DMSO 0.25%) 40 min. before session start according to their weight. These daily injections of vehicle required an adjustment of the water drop size for each mouse to keep them motivated to do 150 trials per session. On the reversal day and the 2 following days, for experimental mice, CNO was diluted in the vehicle solution and delivered at a concentration of 3 mg/kg. In both reversal learning tasks used, we ensured that mice could correctly perform the task in at least 3 consecutive days before reversing the odour-outcome contingency for the first time. In the reversal day, mice started the session as before and the contingencies were reversed at trial 50 in the hM4D experiment and between the 32th and the 100th trial (73 ± 12, mean ± SD) in the GCaMP6s experiments. One SERT-Cre mouse was excluded from the hM4D analysis for not showing differential lick rate in 1.5 s starting at US delivery between odour 1 or 2 (rewards) and odours 3 and 4 (nothing). Two mice were excluded from the GCaMP6s data analysis for bad fibre placement assessed after histology analysis (more than 400 μm away from the infection area): one SERT-Cre and one TH-Cre mouse. Additionally, another SERT-Cre mouse was discarded from the reversal data analysis because of experimental problems with the fibre during the reversal session. In 4 SERT-Cre mice and in all TH-Cre mice, at 5 to 6 days after reversal, we introduced surprise trials during the task. These surprise trials represented approximately 20% of the total number of trials in a session during which no odour cue was presented: the typical white noise of the foreperiod was immediately followed by one of the four possible outcomes, randomly selected (11 ± 4 surprise vs 44 ± 8 cued trials per session, mean ± SD). To analyse these data 4 sessions with predicted and unpredicted outcomes were pooled together for each mouse. All GCaMP6s experiments were performed within the limit of 1 month since viral injection date to avoid cell death due to overexpression of GCaMP6s in neurons.

**Fibre photometry setup** - The dual-color fibre photometry acquisition setup consists of a 3-stage tabletop black case containing optical components (filters, dichroic mirrors, collimator), two light sources for excitation and two photomultiplier tubes (PMTs) for acquisition of fluorescence of a green (GCaMP6s) and of a red (tdTomato) fluorophore. We used a 473 nm (maximum power: 30mW) and a 561 nm (maximum power: 100mW) diode-pumped solid-state laser (both from Crystalaser) for excitation of GCaMP6s and of tdTomato respectively. Beamsplitters (BS007, Thorlabs) and photodiodes (SM1PD1A, Thorlabs) were used to monitor the output of each laser. The laser beams were attenuated with absorptive neutral density filters (Thorlabs) and each was aligned to one of the two entrances of the 3-stage tabletop black case (Doric Lenses). At the corresponding entrances the excitation filters used were 473 nm (LD01-473/10-25 Semrock) and 561nm (LL02-561- 25 Semrock). Inside the black case 3 interchangeable/stackable cubes (Doric Lenses) with dichroic mirrors were used: one to separate the 473nm excitation light from longer wavelengths (Chroma T495LP), one to collect the emission light of GCaMP6s (FF552- Di02-25x36 Semrock), and one to separate the 561nm excitation light from tdTomato’s fluorescence (Di01-R561-25x36). A collimator (F = 12 mm NA = 0.50, Doric Lenses) focused the laser beams in a single multimode silica optical fibre with 300 μm core and 0.22 NA (MFP_300/330/900-0.22_2.5m-FC_CM3, Doric Lenses), which was used for transmission of all excitation and emission wavelengths. The 3-stage tabletop black case had two exits, one for each fluorophore emission, at which we placed the corresponding emission filters (Chroma ET525/50m for GCaMP6s and Semrock LP02-568RS-25 for tdTomato), and convergent lenses (F = 40 mm and F = 50 mm, Thorlabs) before the photodetectors (photomultiplier tube module H7422-02, Hamamatsu Photonics). The output signals of the PMTs were amplified by a preamplifier (C7319, Hamamatsu) and acquired in a Micro1401-3 unit at 5000 Hz and visualized in Spike2 software (Cambridge Electronic Design).

Light power at the tip of the patchcord fibre was 200 μW for each wavelength (473nm and 561 nm) for all experiments (measured before each experiment with a powermeter PM130D, Thorlabs). This patchcord fibre was attached to the fibre cannula each animal had implanted (MFC_300/330-0.22_5mm_RM3_FLT Fibre Polymicro, polymide removed) through a titanium M3 thread receptacle.

**Data Analysis** - All data were analysed in MATLAB. For the hM4D experiment, lick rate was acquired at 1KHz and smoothed using convolution with a Gaussian filter of 50 ms. Mean anticipatory licking was calculated for each trial as the mean lick rate in the period of 500-2800 ms after odour onset subtracted by the mean lick rate in 1s before this period. For each mouse, trials of sessions around reversal were concatenated and a 3 trials smoothing was performed along trials. For each reversed odour and each mouse, the last 50 trials before reversal were fit by a constant function of the form (A+B); the first 200 trials after reversal were fit by an exponential function of the form (A+B*exp(−t/τ)) using *fminsearch* in MATLAB. Conditions for this fitting to be done were: the last 100 trials before reversal had to be statistically different from trial 100-200 after reversal (t-test), the change in licking pattern had to follow the correct trend of the reversal (increase in licking for positive reversals and decrease in licking for negative ones), the time constant obtained had to be larger than 1. Mouse-odour pairs that did not fill this condition were excluded (i.e. odour 4 of mice M#4 and M#5). Time constants were grouped according to the type of reversal and genotype with drug treatment and compared using One-way-ANOVA. Then, for each SERT-Cre mouse, the time constant of the reversal with vehicle was subtracted from the reversal with CNO. The same was done for WT mice, but subtracting the time constant of reversal 2 from that of reversal 1 (since CNO was delivered in both). *t*-tests were used to evaluate if these differences had means significantly different from zero.

Fluorescence data were downsampled to 1 kHz and smoothed using convolution with a gaussian filter of 100 ms. For each trial, the relative change in fluorescence, ΔF/F_0_ = (F-F_0_)/F_0_, was calculated by considering F_0_ the mean fluorescence during 1 s before the odour presentation for both the red and the green channels ([ΔF/F_0_]_GREEN_ and [ΔF/F_0_]_RED_). For each session of each mouse, the distribution of green to red values of ΔF/F_0_was fitted by the sum of two Gaussians along the red channel and the crossing point between these 2 Gaussians was used as a boundary (excluding the first and last 1000 ms of each trial because of filtering artefacts). All values of [ΔF/F_0_]_RED_ below this boundary were used, together with the corresponding [ΔF/F_0_]_GREEN_, to fit a linear regression line. Then, for each trial we corrected the green ΔF/F_0_ values using the parameters (*a* - slope; *b* - offset) obtained with the regression model of that mouse in that session: [ΔF/F_0_]_GREEN___corr_ = [ΔF/F_0_]_GREEN_ - *a\**[ΔF/F_0_]_RED_ - *b*.

Behavioural data was organized as a function of US type and divided in CS and US responses. [ΔF/F_0_]_GREEN___corr_ US responses were normalized by subtracting the mean [ΔF/F_0_]_GREEN___corr_ of the interval of 1 s before US onset. The CS or US response was considered the mean of the signal during 1.5 s after CS or US onset respectively. For each mouse, all CS and US responses were z-scored in the expert phase to compare the amplitudes of responses to the different events. Analysis of US response across time was done by z-scoring all US responses of each mouse across days for each US type. Statistical analysis was done by comparing each day with pre-reversal days -1 and -2. For each mouse, mean amplitude of response to each US on the reversal day was also compared to the day before reversal. For analysis of surprise trials, four days of each mouse were pooled together due to the small number of surprise trials of each US type in each session.

For analysis of CS response time course during reversal, each mean amplitude change across trials was fitted by an exponential with maximum time constant of 225 trials (minimum number of trials after reversal for any US type of any mouse). The same criteria and parameters used for the hM4D experiments were used here. Time constants for mouse-odour pairs were pooled together in pairs (odours 1 and 2, odours 3 and 4) which correspond to the negative and positive reversals respectively.

**Immunohistochemistry and anatomical verification** - Mice were deeply anesthetized with pentobarbital (Eutasil, CEVA Sante Animale), exsanguinated transcardially with cold saline and perfused with 4% paraformaldehyde (P6148, Sigma-Aldrich). Coronal sections (40 μm) were cut with a vibratome and used for immunohistochemistry. For SERT-Cre mice used in expression specificity analysis, anti-5-HT (36 h incubation with rabbit anti-5- HT antibody 1:2000, Immunostar, followed by 2h incubation with Alexa Fluor 594 goat anti-rabbit 1:1000, Life Technologies) and anti-GFP immunostainning (15 h incubation with mouse anti-GFP antibody 1:1000, Life Technologies, followed by 2h incubation with Alexa Fluor 488 goat anti-mouse 1:1000, Life Technologies) were performed sequentially. For SERT-Cre mice used in behavioural experiments, anti-GFP immunostainning was performed (15h incubation with rabbit polyclonal anti-GFP antibody 1:1000, Life Technologies, followed by 2h incubation with Alexa Fluor 488 goat anti-rabbit 1:1000, Life Technologies).

For TH-Cre mice, anti-GFP (15h incubation with rabbit polyclonal anti-GFP antibody 1:1000, Life Technologies, followed by 2h incubation with Alexa Fluor 488 goat antirabbit 1:1000, Life Technologies) and anti-TH immunostainning (15h incubation with mouse monoclonal anti-TH antibody 1:5000, Immunostar, followed by 2h incubation with Alexa Fluor 647 goat anti-mouse, 1:1000, Life Technologies) were performed sequentially. To quantify the specificity of GCaMP6s expression in 5-HT neurons of SERT-Cre mice we used a confocal microscope (Zeiss LSM 710, Zeiss) with a 20X objective (optical slice thickness of 1.8 μm) to acquire z-stacks of three slices around the center of infection. Images for DAPI, GFP and Alexa Fluor 592 were acquired and quantification of cells expressing GCaMP6s and of cells stained with 5-HT antibody were quantified in a 200 × 200 μm window in the center of the DRN. The same was done for quantification of specificity in DA neurons of TH-Cre mice, but acquising Alexa Fluor 647 instead of 592 and taking the 200 × 200 |im window on the infection side. To evaluate fibre location in relation to infection, images for DAPI, YFP or GFP and tdTomato were acquired with an upright fluorescence scanning microscope (Axio Imager M2, Zeiss) equipped with a digital CCD camera (AxioCam MRm, Zeiss) with a 10X objective. The location of the fibre tip was determined by the most anterior brain damage made by the optical fibre subtracted by its radius. The center of infection was estimated through visual inspection of slices as the location where there were most infected cells. The distance between the fibre tip location and center of infection was calculated as an anterior-posterior distance which was estimated by comparing each corresponding location in the mouse brain atlas ^49^ To determine the overlap between cells expressing YFP or GCaMP6s and tdTomato in SERT-Cre mice, we used a confocal microscope (Zeiss LSM 710, Zeiss) with a 20X objective (optical slice thickness of 1.8 μm) to image three slices around the center of infection (slices −1, 0 and 1, relative to it). All cell counts were done using the Cell Counter plugin of Fiji.

